# Virally Encoded Connectivity Transgenic Overlay RNA sequencing (VECTORseq) defines projection neurons involved in sensorimotor integration

**DOI:** 10.1101/2021.10.27.465895

**Authors:** Victoria Cheung, Philip Chung, Max Bjorni, Varvara A. Shvareva, Yesenia C. Lopez, Evan H. Feinberg

**Author notes:** These authors contributed equally (Max Bjorni, Varvara A. Shvareva).

## Abstract

Behavior arises from concerted activity throughout the brain. Consequently, a major focus of modern neuroscience is defining the physiology and behavioral roles of projection neurons linking different brain areas. Single-cell RNA sequencing has facilitated these efforts by revealing molecular determinants of cellular physiology and markers that enable genetically targeted perturbations such as optogenetics, but existing methods for sequencing of defined projection populations are low-throughput, painstaking, and costly. We developed a straightforward, multiplexed approach, Virally Encoded Connectivity Transgenic Overlay RNA sequencing (VECTORseq). VECTORseq repurposes commercial retrogradely infecting viruses typically used to express functional transgenes, e.g., recombinases and fluorescent proteins, by treating viral transgene mRNA as barcodes within single-cell datasets. VECTORseq is compatible with different viral families, resolves multiple populations with different projection targets in one sequencing run, and identifies cortical and subcortical excitatory and inhibitory projection populations. Our study provides a roadmap for high-throughput identification of neuronal subtypes based on connectivity.

## Introduction

Functionally and molecularly diverse projection neurons with distinct targets are intermingled within most brain areas. For example, primary visual cortex contains functionally distinct populations that project to higher visual cortical areas, contralateral cortex, and subcortical targets such as striatum, thalamus, and superior colliculus; amygdala contains different projection neurons that target the brainstem, hippocampus, thalamus, olfactory structures, and several cortical areas; and substantia nigra pars reticulata in midbrain harbors subtypes that project to 39 target structures and differ in neurotransmitters released, response tuning, and intrinsic excitability (Antal et al., 2014; Janak and Tye, 2015; Jiang et al., 2003; Kim et al., 2015; Lur et al., 2016; McElvain et al., 2021; Poulin et al., 2016). A major goal of modern neuroscience is to understand the properties and behavioral functions of the myriad projection neuron subtypes in the brain. Single-cell RNA sequencing technologies hold the promise of providing insights into the electrophysiology of each identified neuronal population and molecular markers that could be used for targeted monitoring and manipulation of these cell types during behavior. However, the major challenge in interpreting single-cell RNA sequencing datasets is to link populations of cells identified through RNA expression to their anatomy, connectivity, and circuit properties, a conundrum known as a “correspondence problem” (Lein et al., 2017).

In recent years, a few approaches have been developed for targeted transcriptional profiling of projection populations. Many take advantage of dyes or viruses that are injected into the target structure, internalized at axon terminals, and trafficked retrogradely to label cell bodies in the source structure (Wickersham and Feinberg, 2012). In one approach, retrogradely labeled cells in the source structure are isolated from dissociated tissue on the basis of their fluorescence and then sequenced (Tasic et al., 2018). In another approach, Patch-seq, cells are targeted for intracellular recording before the cellular contents are aspirated and sequenced (Cadwell et al., 2017). Both approaches are laborious, requiring separate rounds for each projection target and population of interest and additional steps for selective isolation of labeled cells, and costly, because each projection target is sequenced separately. A third sequencing-based approach, retroTRAP (translating ribosome affinity purification), which does not provide single-cell resolution, relies on the use of retrogradely delivered recombinases to induce expression of tagged ribosomal subunits (or other mRNA-binding proteins) in projection neurons followed by immunoprecipitation of tagged mRNA from the projection cell type(s) of interest (Ekstrand et al., 2014). Because this method does not resolve single cells, it obscures heterogeneity within projection populations; moreover, it must be performed iteratively on just one projection target at a time. A recently described sequencing-based method, MAPSeq, uses a different approach to trace connectivity anterogradely (Kebschull et al., 2016). A structure of interest is infected with Sindbis virus encoding RNA barcodes that are trafficked into axons and detected by RNA sequencing of target structures. To relate projections to cell types, BARseq combines MAPseq with *in situ* sequencing of starter cells to identify their Sindbis-encoded barcodes and fluorescence *in situ* hybridization (FISH) to assign these starter cells to particular cell types (Xiaoyin Chen et al., 2019). BARseq allows identification of multiple projection populations at once but requires several additional steps and specialized equipment, including for *in situ* sequencing. Moreover, the only published application of BARseq required first using standard single-cell sequencing approaches to identify candidate markers for FISH. Another limitation of this approach is that Sindbis virus is highly toxic and rapidly disrupts cellular transcription, and requires calibration of expression times for every experiment. Thus, there is a pressing need for a straightforward method for multiplexed transcriptional profiling of myriad projection cell types in the transcriptional “ground state" in a single sequencing experiment without specialized additional equipment. Such a method would be transformative because it would lower barriers to access, increase throughput, reduce the numbers of animals used per experiment, expedite experiments, and reduce costs.

We reasoned that retrograde viral methods that have been widely used to selectively label projection populations (*e.g.*, with fluorophores) could be repurposed to deliver mRNA barcodes that are directly detected in single-cell sequencing datasets. This would enable multiplexed identification of many different projection cell types in one single-cell sequencing experiment without additional equipment. Here we describe a method for doing so, Virally Encoded Connectivity Transgenic Overlay RNA sequencing (VECTORseq). We show that virally encoded transcripts delivered via retrograde infection are robustly detected by single-cell sequencing, readily distinguished from closely related isoforms, and found selectively in the expected populations in primary visual cortex. We then apply VECTORseq to multiple subcortical structures involved in sensorimotor integration. We show that VECTORseq enables multiplexed identification of both known and novel projection populations in the transcriptional “ground state.” Our study thus establishes a straightforward, high-throughput method to transcriptionally profile projection populations and identifies new subcortical cell types involved in sensorimotor integration.

## Results

VECTORseq is based on the idea that widely used retrogradely infecting viruses express transgenes such as recombinases, fluorophores, and optogenetic molecules whose mRNA could be treated as projection-based RNA barcodes to overlay anatomy on single-cell sequencing data. For example, if we inject structure A with a retrogradely infecting virus encoding green fluorescent protein (GFP) and structure B with retrogradely infecting virus encoding Cre recombinase, cells that project to structure A will express *GFP* mRNA, whereas cells that project to structure B will express *Cre* mRNA (Fig. 1). Thus, in a single-cell sequencing dataset of structure C, cells expressing the Cre or GFP mRNA can be identified as projecting to structure A or B.

**Figure 1.**
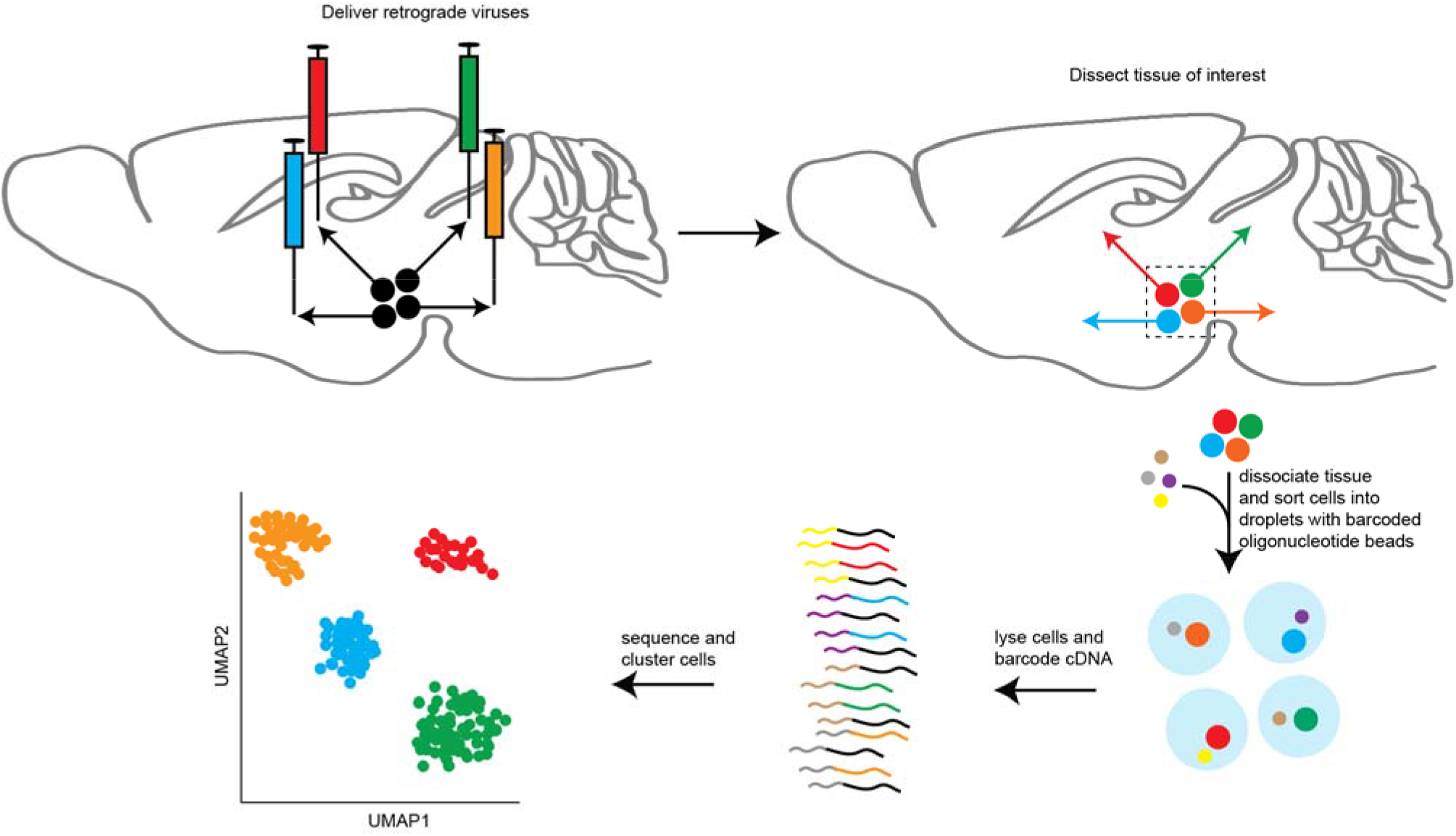
Schematic of VECTORseq approach. Injecting retrogradely infecting viruses (colored syringes) into brain structures (sagittal section of mouse brain in this schematic) targeted by different projection neurons (directed arrows) from a single structure of origin will label each with unique virally encoded RNA barcodes. Tissue is then dissociated, cells (large dots) are sorted into droplets with beads (small dots) coated in barcoded oligonucleotides, tagging all RNA from each cell with a cell-specific barcode, including the viral transgene (barcoded barcode). Once RNA from each cell is identified according to the cell-specific barcodes and cells are clustered, expression of viral barcodes can be overlaid to assign each cell-type cluster to its projection target and functional properties.

### Validation of VECTORseq in primary visual cortex

We first tested the feasibility of the VECTORseq approach on projection populations in primary visual cortex (V1). These populations offered an appealing benchmark because they had previously been transcriptionally profiled using fluorescence to isolate retrogradely labeled cells (Tasic et al., 2018). Our goal was to define transcriptional “ground states,” and the widely used adeno-associated virus (AAV) is thought to express transgenes without perturbing cellular physiology (Haggerty et al., 2020). In addition, commercial sources (*e.g.,* Addgene and university vector cores) offer retrogradely infecting AAV carrying diverse transgenes (*e.g.*, *GFP, tdTomato, Cre, FLP*) that could be treated as multiplexable barcodes. Therefore, in five male mice, we injected off-the-shelf AAV retrograde (AAVrg) encoding different transgenes into three major V1 projection targets: *mCherry* and *Cre* separated by an internal ribosomal entry site (IRES) under control of the *EF1*α promoter (*EF1*α*-mCherry-IRES-Cre*) in left striatum; FLPo recombinase under control of the EF1α promoter (*EF1*α*-FLPo*) in left superior colliculus (SC); and Dre recombinase under control of a human synapsin promoter element (*hSyn-Dre*) in right (contralateral) V1 (Fig. 2A) (Kim et al., 2015; Lur et al., 2016; Tasic et al., 2018; Tervo et al., 2016). To be able to visualize the injection sites for viruses encoding transgenes whose protein products are not fluorescent (FLPo and Dre), we included *AAV1-hSyn-TurboRFP*; because AAV1 can also traffic retrogradely in a dose-dependent manner, we heavily diluted this virus to reduce retrograde labeling (Tervo et al., 2016). In addition, we injected left V1 with AAV1 encoding a Cre-dependent form of the red fluorescent protein tdTomato under control of the synthetic CAG promoter (*CAG-FLEX-tdTomato*) and AAV1 encoding a FLP-dependent form of yellow fluorescent protein (*EF1*α*-fDIO-EYFP*) to provide fiducials for microdissection of V1 before dissociation and, if necessary, signal amplification for the retrogradely delivered *mCherry-IRES-Cre* and *FLPo*, respectively, in the sequencing dataset (Fig. 2A). After waiting 3 weeks for retrograde trafficking of viruses and transgene expression, we dissected left V1, dissociated tissue into single cells, and processed cell suspensions for single-cell sequencing using the 10x Chromium system (Fig. 2A). We used the Chromium 5’ kit because many commercial AAVs incorporate the Woodchuck Hepatitis Virus Posttranscriptional Regulatory Element (WPRE) in the 3’ UTR to boost transgene expression (Wang et al., 2016). Sequencing was then performed using standard Illumina paired-end sequencing on their NextSeq platform. For this pilot experiment, sequencing depth was relatively shallow and relatively few cells were sequenced.

**Figure 2:**
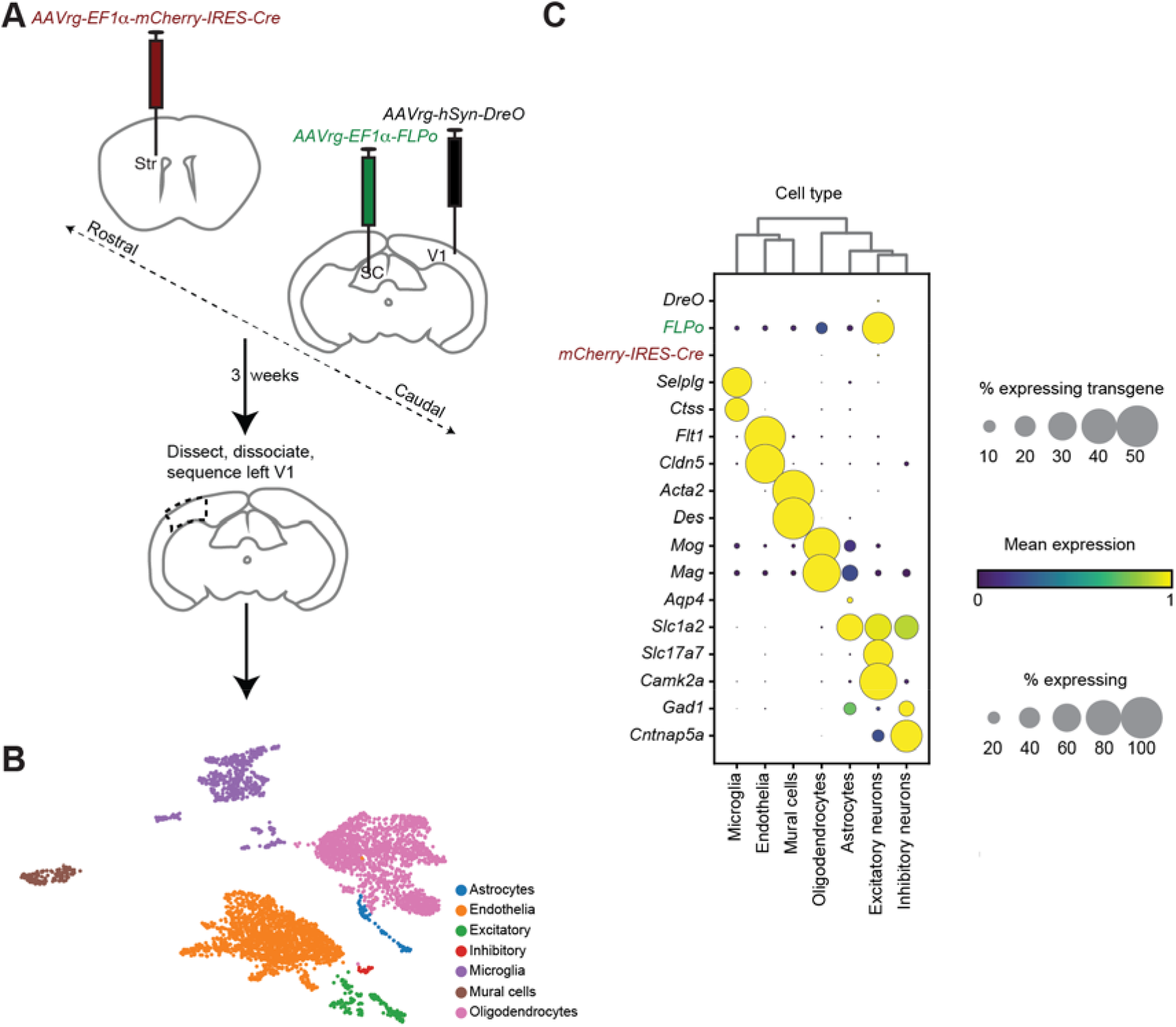
VECTORseq of V1 projection neurons. **A**. Schematic of experimental approach. Retrograde viruses were injected into three major projection targets of left V1: ipsilateral (left) SC, ipsilateral (left) striatum, and contralateral (right) V1. To mark injection sites in SC and contralateral SC, dilute *AAV1-hSyn-TurboRFP* was co-injected (not illustrated). To provide a fiducial for microdissection and signal amplification for sequencing, if needed, left V1 was also injected with *AAV1-CAG-FLEX-tdTomato* and *AAV1-EF1*α*-fDIO-EYFP* (not illustrated). 3 weeks later, left V1 was dissected, cells were dissociated, and single-cell sequencing was performed. **B**. UMAP of V1 sequencing data illustrating different major cell types in this dataset. **C**. Enriched and differentially expressed genes in major cortical cell types and distribution of retrograde viral transgenes. Note the different scales for transgene and endogenous gene expression.

Analyses of the sequencing dataset revealed 21,702,272 reads corresponding to 4,167 cells. We first determined whether the retrograde transgenes were detected. We observed 15 reads aligned to *mCherry-IRES-Cre*; 8,007 reads aligned to *FLPo*; and 1 read aligned to *Dre*. Within cells expressing the viral transgenes, the expression level was as high as that of many commonly used marker genes (Table 1). In addition, despite having attempted to reduce retrograde trafficking of AAV1-hSyn-TurboRFP used to mark the injection sites by diluting the virus, we detected 114 reads of the *TurboRFP* construct injected into SC and contralateral V1. To determine the specificity of VECTORseq, we included in our library the sequences of the injected *mCherry-IRES-Cre* and another *Cre* isoform found in commercial AAV and other viruses; these isoforms have codon substitutions that render them 74% identical at the RNA sequence level (Supplementary Figure 1). Whereas 15 reads aligned to *mCherry-IRES-Cre*, no reads were aligned to the other *Cre* sequence. Thus, VECTORseq is sufficiently sensitive to detect retrograde transcripts and sufficiently specific to discriminate homologous transgenes in single-cell datasets. Although the robust detection of the Cre transgene obviated the need for the FLEX and fDIO reporters for signal amplification, we detected abundant *tdTomato* and *EYFP* reads (33,983 reads in 769 cells and 10,045 reads in 350 cells, respectively). This widespread *tdTomato* and *EYFP* detection was likely due to leaky antisense transcription or recombination in bacteria during plasmid production (Fischer et al., 2019).

**Table 1:** Transgene detection in V1 sequencing dataset. First column lists transgenes and endogenous genes for comparison. Second column indicates the number of cells in which each transgene or endogenous gene was detected. Third column indicates total number of reads corresponding to transgenes or endogenous genes in V1 sequencing dataset overall. Fourth column indicates relative expression of transgenes and common marker genes. To control for differences in abundance of different cell types, values denote the mean percentage of reads +/-standard deviation corresponding to a given marker or transgene in cells positive for that marker or transgene.

| Transgenes | Number of expressing cells | Number of reads | Abundance in expressing cells |
| --- | --- | --- | --- |
| *Dre* | 1 | 1 | 0.007 |
| *FLPo* | 216 | 8,007 | 0.43 +/- 1.38 |
| *mCherry-IRES-Cre* | 3 | 15 | 0.70 +/- 0.05 |
| *turboRFP* | 35 | 114 | 0.05 +/- 0.06 |
| *EYFP* | 350 | 10,045 | 0.85 +/- 2.73 |
| *tdTomato* | 769 | 33,982 | 0.96 +/- 3.38 |
| **Endogenous Genes** |  |  |  |
| *Snap25* | 313 | 3,522 | 0.14 +/- 0.14 |
| *Rbfox3* | 138 | 459 | 0.04 +/- 0.03 |
| *Slc17a6* | 72 | 152 | 0.04 +/- 0.03 |
| *Camk2a* | 169 | 912 | 0.07 +/- 0.06 |
| *Gad1* | 44 | 265 | 0.04 +/- 0.05 |
| *Gad2* | 10 | 176 | 0.07 +/- 0.04 |
| *Mog* | 1,482 | 19,362 | 0.21 +/- 0.12 |
| *Flt1* | 1,654 | 20,198 | 0.27 +/- 0.21 |

We used the graph-based Leiden algorithm to cluster cells and Uniform Manifold Approximation and Projection (UMAP) to visualize and annotate clusters based on expression of known marker genes, separating neurons, endothelia, and different classes of glial cells (Fig. 2B) (Chamling et al., 2021; Chen et al., 2020; Hammond et al., 2019; Hasel et al., 2017; He et al., 2016; Traag et al., 2019; Yao et al., 2021). We then subclustered neurons into excitatory and inhibitory subpopulations and overlaid expression of the viral transgenes. We predicted that the retrograde transgenes should be in neurons rather than non-neuronal cells. Moreover, because the aforementioned studies that used sorting and sequencing of fluorescent retrogradely labeled cortical neurons that project to SC, striatum, and contralateral cortex found that these cells included a variety of mostly excitatory neurons found in cortical layers 2-6, we predicted that the transgenes would be enriched in the cluster containing excitatory subtypes (Tasic et al., 2018). Consistent with these predictions, all retrograde transgenes were enriched in the cluster of excitatory neurons (Fig. 2C). These data indicate the feasibility of the VECTORseq approach— transgenes delivered by retrogradely infecting viruses can be detected in single-cell sequencing datasets and their expression is found in the expected cell types.

### Application of VECTORseq to superior colliculus projection populations

To determine whether VECTORseq is applicable in subcortical structures and to attempt to identify new projection populations with it, we next investigated superior colliculus (SC), which harbors a mixture of known and uncharacterized projection populations. We targeted two SC cell types that innervate brainstem on opposite sides: neurons that control orienting movements and innervate the contralateral paramedian pontine reticular formation (PPRF, also known as parabducens) and neurons that drive avoidance responses and innervate the ipsilateral cuneiform nucleus (CnF) (Dean et al., 1989; Sahibzada et al., 1986). A previous study had labeled both populations with retrogradely infecting lentiviruses, but we were unable to identify a commercial source for these retrograde lentiviruses (Isa et al., 2020). Therefore, we attempted to use AAVrg instead, injecting right PPRF with *AAVrg-CAG-GFP* and left CnF with *AAVrg-CAG-tdTomato* (Fig. 3A). Additionally, we targeted SC projections to the lateral posterior nucleus of the thalamus (LP), the rodent homolog of primate pulvinar. Anterograde tracing from SC labels multiple subdivisions of LP, including LPLR and LPMR, and studies in primates have found projections from superficial and deep layers of SC project to the homologous regions of pulvinar (Benevento and Fallon, 1975; Gale and Murphy, 2014; Gharaei et al., 2020; Homman-Ludiye and Bourne, 2019). A serendipitously discovered Cre transgenic line, *Ntsr1-GN209*, labels a cell type in superficial SC, wide-field (WF) cells, that projects to LPLR and is implicated in visual processing and fear responses (Gale and Murphy, 2014; Gerfen et al., 2013). In the Allen *in situ* database, endogenous *Ntsr1* does not appear to be expressed in superficial SC, including WF cells, suggesting that this transgenic line functions as an enhancer trap.

**Figure 3.**
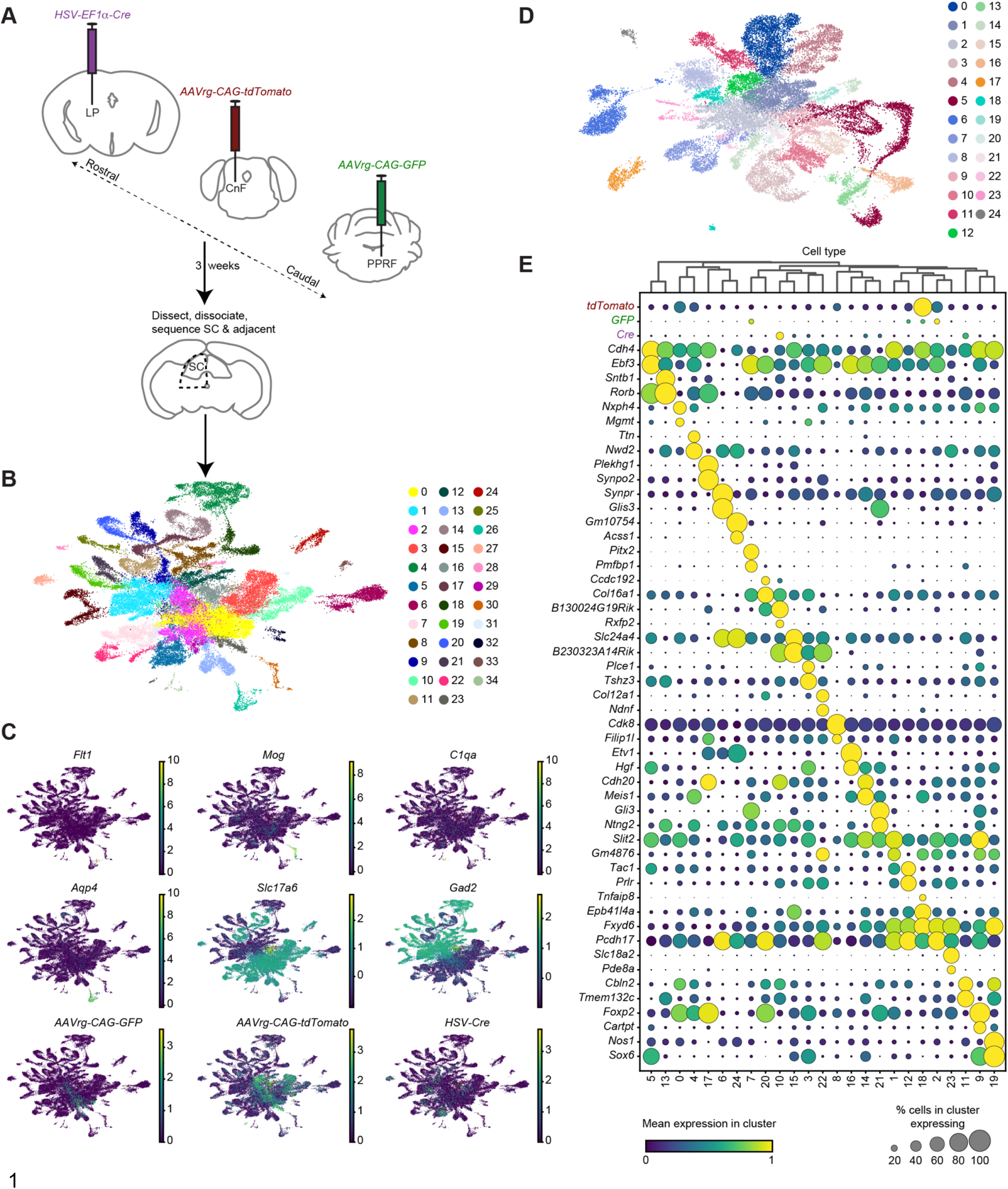
VECTORseq of superior colliculus. **A.** Schematic of experimental approach. Retrograde viruses were injected into left LP, left CnF, and right PPRF. 3 weeks later, left SC was dissected, nuclei were isolated, stained, and sorted according to *NeuN/Rbfox3* expression, and single-cell sequencing was performed. **B**. UMAP of SC sequencing data and clustering. **C.** Overlay of markers for different cell types (*Flt1*, endothelia; *Mog*, oligodendrocytes; *C1qa*, microglia; *Aqp4*, astrocytes, *Slc17a6*, excitatory neurons; *Gad2*, inhibitory neurons) shows enrichment of neurons in this dataset and presence of transgenes in neuronal clusters. **D.** UMAP of SC excitatory (*Slc17a6*^+^) neurons. **E.** Enriched and differentially expressed genes in major SC excitatory cell types and distribution of retrograde viral transgenes.

Nevertheless, this Cre line has become a popular tool for studies of the involvement of SC and LP in visual processing and behavior (Gale and Murphy, 2014, 2016; Hoy et al., 2019; Reinhard et al., 2019; Sans-Dublanc et al., 2021). However, the projection from deep SC to LPMR remains mysterious, with its molecular and functional properties unknown. We therefore targeted LP in hopes of transcriptionally profiling both WF cells and this undefined LPMR- projecting population. Because a recent study used another retrograde virus that is reported to preserve cellular ground states, herpes simplex 1 (HSV-1), to retrogradely infect WF cells, we decided to test the applicability of VECTORseq to other viral families (Neve et al., 2018; Reinhard et al., 2019; Verlengia et al., 2017). We injected HSV-1 encoding Cre near the border between LPLR and LPMR to label the different SC neurons projecting to each structure (Fig. 3A).

Once again, we waited 3 weeks for retrograde trafficking of viruses and transgene expression before dissociating SC and surrounding structures. We dissected the dorsal midbrain, containing SC and adjacent structures, such as portions of periaqueductal gray and inferior colliculus, and pooled tissue from 4 mice. Because sequencing nuclei rather than whole cells has become a popular technique, as has enriching for neuronal nuclei using flow cytometry, we determined the compatibility of VECTORseq with nuclear sequencing by isolating nuclei expressing the neuronal marker *NeuN/Rbfox3* (Krishnaswami et al., 2016). We generated single-nucleus libraries using 10X Chromium 5’ kits and performed Illumina sequencing using the NovaSeq platform for greater depth. These analyses revealed 628,122,200 reads corresponding to 54,537 cells. We observed 170,025 reads (in 11,377 cells) aligned to *AAVrg-CAG-tdTomato*, 5,270 reads (in 1,531 cells) aligned to *AAVrg-CAG-GFP*, and 15,861 reads (in 3,117 cells) aligned to *HSV-Cre*, showing the compatibility of VECTORseq with single-nucleus sequencing approaches and with viruses other than AAV. As in cortex, these viral transgenes were highly expressed at levels comparable to those of commonly used cell-type markers such as *Rbfox3, Slc17a6*, and *Gad1* (Table 2). Importantly, in this experiment we once again detected 15,861 reads corresponding to the injected *Cre* isoform, and zero reads that aligned to the other *Cre* isoform in our library (*i.e*., *mCherry-IRES-Cre,* which was used in the V1 sequencing experiment), further demonstrating the specificity of VECTORseq in discriminating different transgenes.

**Table 2:** Transgene detection in SC sequencing dataset. First column lists transgenes and endogenous genes for comparison. Second column indicates the number of cells in which each transgene or endogenous gene was detected. Third column indicates total number of reads corresponding to transgenes or endogenous genes in SC sequencing dataset overall. Fourth column indicates relative expression of transgenes and common marker genes. As a control for differences in abundance of different cell types, values denote the mean percentage of reads +/-standard deviation corresponding to a given marker or transgene in cells positive for that marker or transgene.

| Transgenes | Number of expressing cells | Number of reads | Abundance in expressing cells |
| --- | --- | --- | --- |
| *Cre* | 3,117 | 15,861 | 0.05 +/- 0.13 |
| *GFP* | 1,531 | 5,270 | 0.03 +/- 0.04 |
| *tdTomato* | 11,377 | 170,025 | 0.70 +/- 0.05 |
| **Endogenous Genes** |  |  |  |
| *Snap25* | 53,717 | 617,837 | 0.10 +/- 0.05 |
| *Rbfox3* | 45,279 | 155,127 | 0.03 +/- 0.02 |
| *Slc17a6* | 31,559 | 126,995 | 0.03 +/- 0.02 |
| *Camk2a* | 39,291 | 121,268 | 0.03 +/- 0.03 |
| *Gad1* | 26,957 | 116,424 | 0.04 +/- 0.04 |
| *Gad2* | 28,342 | 159,007 | 0.05 +/- 0.04 |
| *Mog* | 2,842 | 4,856 | 0.02 +/- 0.04 |
| *Flt1* | 86 | 469 | 0.07 +/- 0.13 |

We then performed graph-based clustering of the sequencing dataset to identify different cell types (Fig. 3B, C). The overwhelming majority (98.54%, 53,740/54,537) of the profiled nuclei were neuronal, indicating that the enrichment was successful. We next subclustered excitatory neurons (60.8% of all neurons, 32,658/53,740) and overlaid the expression of viral transgenes on these clusters (Fig. 3D, E). Within clusters, transgene-positive and transgene-negative cells were interspersed, suggesting that the viruses did not perturb endogenous gene expression or the cellular ground state in such a way that virus-expressing cells segregated within gene expression space (Supplementary Fig. 2). We then examined *GFP^+^* cells. *GFP* reads were most prevalent in two clusters. One of these, cluster 7, expressed *Pitx2*, a marker previously shown to label deep SC neurons that drive orienting movements and project to contralateral PPRF and zona incerta (Masullo et al., 2019; Xie et al., 2021). In addition, this cluster expressed markers such as *Pmfbp1* with expression patterns in deep SC similar to that of *Pitx2* in the Allen *in situ* atlas. To confirm that these markers were expressed in the PPRF-projecting population, we injected PPRF with AAVrg encoding *Cre* and SC with *AAV1-FLEX-tdTomato* (to provide signal amplification if *Cre* expression were too weak to detect by *in situ*, which was not the case), waited 3 weeks, and used RNAscope fluorescent *in situ* hybridization (FISH) to determine whether *Cre^+^* cells expressed *Pitx2* or *Pmfbp1* (Fig. 4A) (Wang et al., 2012). 120 of 145 *Cre*^+^ cells (83%, n = 3 animals) also expressed *Pitx2* (Fig. 4A). We next examined *Pmfbp1*. In the sequencing dataset, *Pmfbp1* was detected in a smaller fraction of the cells in this cluster than was *Pitx2*, suggesting that it is expressed at lower levels and leading us to predict it would be detected in fewer *Cre*^+^ cells. Consistent with this prediction, a smaller fraction of *Cre*^+^ cells were *Pmfbp1^+^* (33/67, 49%, n = 4 animals) by FISH (Fig. 4B). To confirm the specificity of *Pmfbp1* as a marker for this population, we examined its expression in a projection population that the sequencing dataset suggested expressed minimal amounts of *Pmfbp1*. We injected *AAVrg-Cre* into LP and used RNAscope to measure *Pmfbp1* co-expression in *Cre*^+^ cells (Fig. 4C). Only 21 of 160 *Cre*^+^ cells were also *Pmfbp1^+^* (13%, n = 5 animals), indicating that *Pmfbp1* is specific to PPRF-projecting cells (p < 0.0001, chi-square test) (Fig. 4C). This result confirmed that *Pmfbp1* is a new marker for this population of deep SC neurons that also express *Pitx2*. Thus, VECTORseq was able to identify a known subcortical projection population and identify new markers for it, confirming the sensitivity and specificity of this approach.

**Figure 4.**
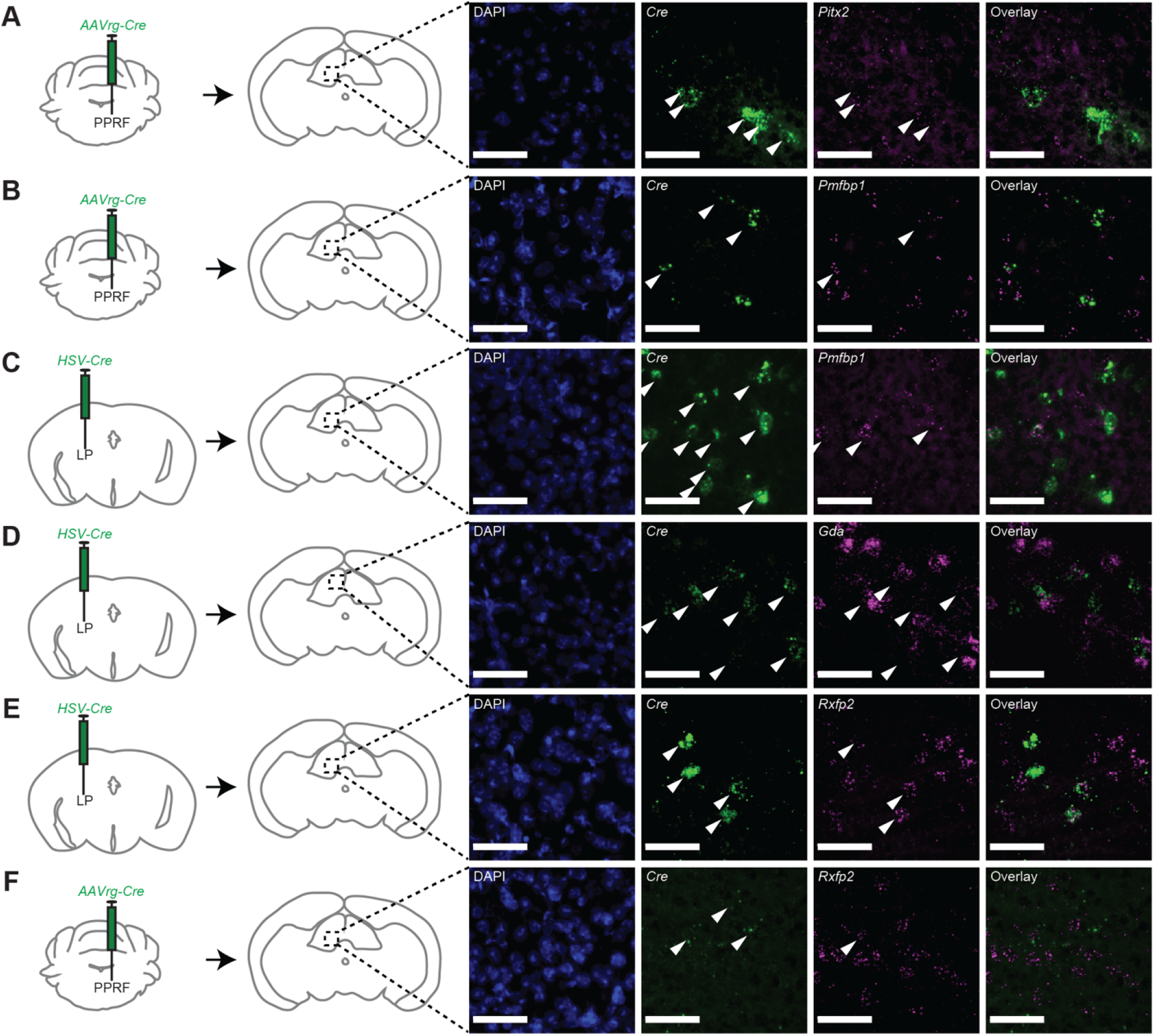
Analysis of candidate marker gene expression in different SC projection populations. **A.** Left, schematic of experiment. *AAVrg-Cre* was injected in right PPRF. 3 weeks later, mice were perfused and RNAscope was performed on the deep layers of left SC. Right, representative images. Middle left image shows expression of *Cre*. Arrowheads indicate *Cre^+^* cells. Middle right image shows *Pitx2* expression. Arrowheads indicate *Cre^+^* cells thatare also *Pitx2*^+^. **B, C**. As in A but for *Pmfbp1* expression in PPRF-projecting (B) and LP-projecting (C) neurons in deep SC. **D**. As in A-C but for *Gda* expression in LP-projecting neurons in superficial SC. **E, F**. As in A-D but for *Rxfp2* expression in LP-projecting (E) and PPRF-projecting (F) neurons in deep SC. Scale bars, 50 μm.

We were surprised to detect *tdTomato* in virtually every neuronal population. This did not seem to be due to sequencing errors, because our libraries included negative control transgene sequences that were not injected, such as *mCherry-IRES-Cre*, that were not detected by sequencing. This led us to wonder whether the retrograde labeling was promiscuous or had spilled over from CnF, which is near SC. Therefore, we examined an identically injected cohort to the cohort that was sequenced. Upon examining SC histologically, we saw sparse tdTomato-labeled fibers but no tdTomato-expressing cells, whereas GFP-labeled, PPRF-projecting cells were abundant (Supplementary Fig. 3A, B). This suggested that retrograde labeling was not promiscuous in SC nor that there was considerable spillover from the injection site. In contrast, the injection site in CnF was extremely brightly labeled (Supplementary Fig. 3C, D). This area abuts SC and was included in our microdissection. This led us to hypothesize that the presence of abundant *tdTomato* reads in every cell cluster was due to contamination by ambient RNA released during nuclei isolation, a common confound, from these extremely strong expressing cells in CnF and surrounding tissue (Alvarez et al., 2020; Yang et al., 2020). When ambient RNA contamination is prevalent, highly expressed genes are found in many populations as the RNA becomes distributed throughout the nuclei suspension (Yang et al., 2020). Therefore, we predicted that *tdTomato* would be equally likely to be detected in neurons and non-neuronal cells. In contrast, if ubiquitous *tdTomato* detection in neuronal populations were due to retrograde infection rather than ambient RNA contamination, *tdTomato* should be found more frequently in neurons than in non-neuronal cells. Consistent with our hypothesis, 1.47% (797/54267) of the total nuclei in our dataset and 1.34% (153/11377) of the *tdTomato*^+^ nuclei were non-neuronal, suggesting that there was not a significant difference in the probability of detecting *tdTomato* in neuronal and non-neuronal cells (p = 0.34, chi-square test). In contrast, the other retrograde transgenes were significantly more likely to be detected in neurons than would be expected by chance: only 0.65% of *GFP*^+^ nuclei were non-neuronal (10/1531, p = 0.009, chi-square test), and only 0.70% of *Cre*^+^ nuclei were non-neuronal (22/3117, p = 0.005, chi-square test). Thus, we concluded that the ubiquity of *tdTomato* reads is due to non-specific contamination by ambient RNA from extremely highly expressing cells in the injection site (CnF) that were included in the dissection. We did not further analyze *tdTomato^+^* populations.

We then analyzed the distribution of *HSV-Cre* labeling in SC excitatory neurons, observing it was most prominent in two populations, clusters 10 and 11 (Fig. 3E). Cluster 11 expressed markers such as *Tmem132c*, *Cbln2*, *Trhde*, and *Gda*, all of which localized to a thin lamina in the stratum opticum, where LPLR-projecting WF cells are found, in the Allen *in situ* dataset; one of these markers, *Cbln2*, was recently shown to be a marker for WF cells (Xie et al., 2021) (Fig. 3E). We used RNAscope to determine whether LP-projecting neurons in superficial SC express *Gda*, which appeared strongly expressed and specific within SC to the stratum opticum in the Allen *in situ* atlas (Fig. 4D). Of 353 retrogradely labeled *Cre^+^* cells in superficial SC, 307 (87%, n = 5 animals) also expressed *Gda*, confirming that it is a marker for WF cells (Fig. 4D). We then analyzed the previously unknown LPMR-projecting population in intermediate and deep SC. This population expressed relatively few unique markers, including the specific but not highly expressed *Rxfp2* (Fig. 3E). We used RNAscope to examine expression of *Rxfp2* in the deep and intermediate SC population that projects to LP. We injected *AAVrg-Cre* into LP, *AAV-CAG-FLEX-tdTomato* into SC (as noted previously, for signal amplification if needed), and waited 3 weeks before performing RNAscope (Fig. 4E). *Rxfp2* was detected in 56 of 213 *Cre*^+^ cells (26%, n = 3 animals). We hypothesized this was reflective of low expression overall, because *Rxfp2*, although specific for that population, was also not highly expressed in the RNA sequencing data (Fig. 3E). Therefore, as a specificity control *Rxfp2*, we examined a projection population that the sequencing dataset suggested expressed minimal amounts of *Rxfp2*. We injected *AAVrg-Cre* into contralateral PPRF and used RNAscope to measure *Rxfp2* co-expression in *Cre*^+^ cells (Fig. 4F). Only 1 of 29 *Cre*^+^ cells also expressed *Rxfp2* (3%, n = 4 animals), confirming the specificity of *Rxfp2* as a marker for the LPMR-projecting population in deep SC (p = 0.0065, chi-square test). To determine whether these clusters were identified in their transcriptional “ground states,” we examined gene expression within these clusters in transgene^+^ and transgene^-^ cells. Importantly, endogenous gene expression did not appear systematically skewed in cells expressing either AAVrg or HSV, indicating that these analyses reveal the cellular “ground state” (Supplementary Fig. 2). Thus, VECTORseq is compatible with multiple viral families and can identify both novel and known subcortical cell types in their “ground states,” including a previously undefined population that projects to LP.

### Application of VECTORseq to ventral midbrain inhibitory projection populations

Many projection populations in the brain, especially in subcortical structures, are inhibitory. Therefore, we tested the applicability of VECTORseq to inhibitory projection populations. We focused on ventral midbrain, where diverse inhibitory projection types in neighboring structures such as zona incerta (ZI) and substantia nigra pars reticulata (SNr), among others, innervate targets involved in movement control, such as motor thalamus (VM), the mesencephalic locomotor region (MLR), and superior colliculus (SC) (Antal et al., 2014; Barthó et al., 2002a; Hikosaka and Wurtz, 1983; McElvain et al., 2021; Nagalski et al., 2015; Watson et al., 2014). We injected VM and MLR with *AAVrg-mCherry-IRES-Cre* and *AAVrg-FLPo*, respectively (Fig. 5A). To test whether VECTORseq is compatible with another commonly used retrograde viral family, we injected SC with Canine Adenovirus 2 (Cav-2) encoding *GFP* (Fig. 5A) (Junyent and Kremer, 2015). In addition, we injected *AAV1-FLEX-tdTomato* in SNr to provide a fiducial for dissections (Fig. 5A).

**Figure 5.**
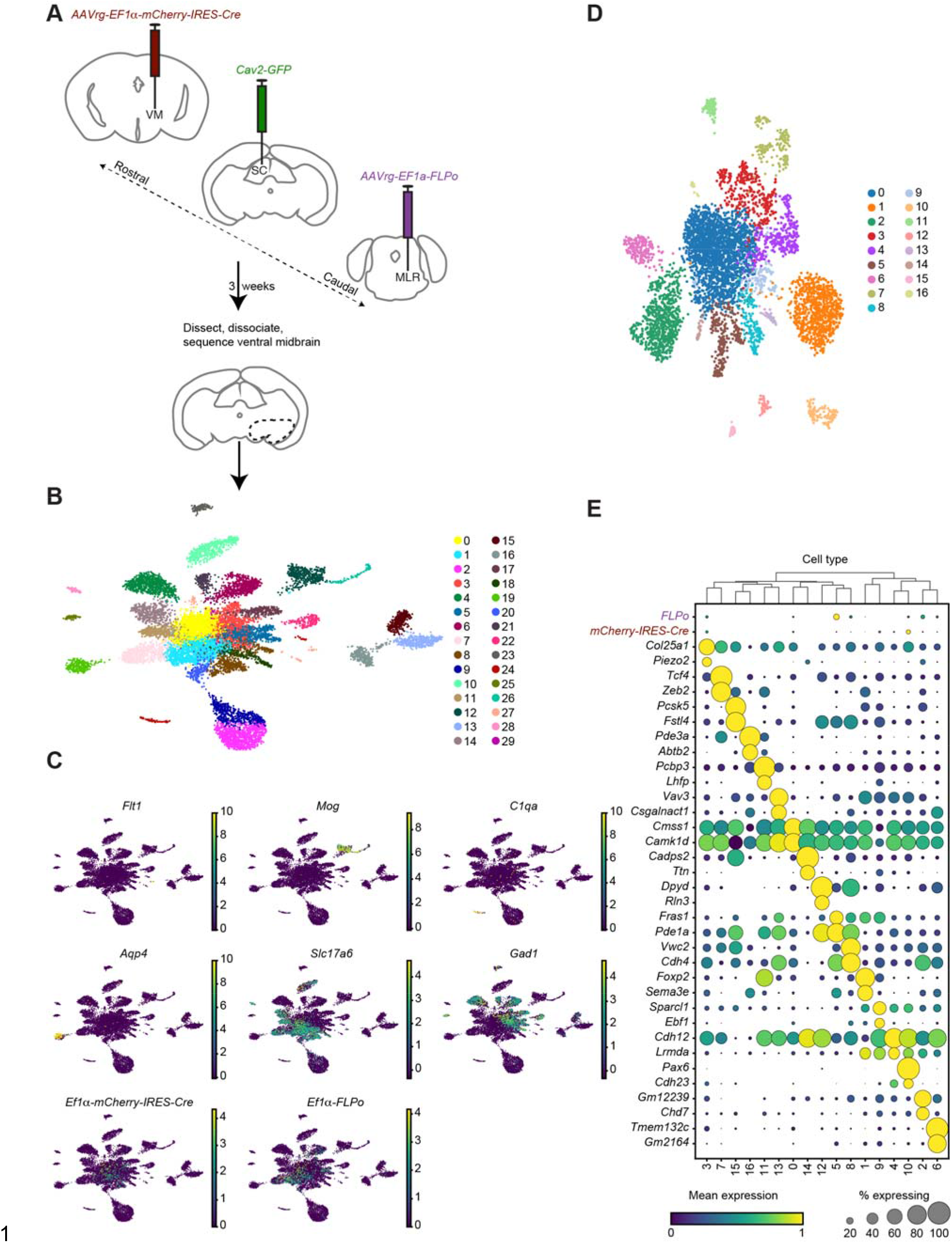
VECTORseq of ventral midbrain inhibitory neurons. **A.** Schematic of experimental approach. Retrograde viruses were injected into right VM, right MLR, and left SC. 3 weeks later, right ventral midbrain was dissected, nuclei were isolated, stained, and sorted, and single-cell sequencing was performed. **B**. UMAP of sequencing data and clustering. **C.** Overlay of markers for different cell types (*Flt1*, endothelia; *Mog*, oligodendrocytes; *C1qa*, microglia; *Aqp4*, astrocytes, *Slc17a6*, excitatory neurons; *Gad1*, inhibitory neurons) shows enrichment of neurons in this dataset and presence of transgenes in neuronal clusters. **D.** UMAP plot of inhibitory (*Gad1^+^/Gad2^+^*) neurons. **E.** Enriched and differentially expressed genes in major ventral midbrain inhibitory cell types and distribution of virally encoded transgenes.

After waiting 3 weeks for viral trafficking and transgene expression, we dissected the portion of the ventral midbrain containing ZI and SNr ipsilateral to the injection sites from 5 mice. Once again, we isolated *NeuN*^+^ nuclei, generated libraries using the 10x Chromium 5’ system, and performed Illumina paired-end sequencing using the NovaSeq platform. Analyses of the sequencing dataset revealed 34,274,388 reads corresponding to 13,412 cells. The *FLP* and *Cre* injected in MLR and VM, respectively, were abundant in the sequencing dataset, at levels comparable to those of commonly examined marker genes (Table 3), whereas the *Cav2-GFP* was not detected. To investigate the lack of *GFP*, we analyzed an identically injected cohort of mice histologically. Because Cav-2 infects both local neurons at the injection site and projections to that site, we examined the injection site in SC. This revealed only a few labeled cells (Supplementary Fig. 4) that fell along the injection track. Thus, the lack of *GFP* reads in the sequencing dataset is likely due to a lack of infection.

**Table 3:** Transgene detection in ventral midbrain sequencing dataset. First column lists transgenes and endogenous genes analyzed. Second column indicates the number of cells in which each transgene or endogenous gene was detected. Third column indicates total number of reads corresponding to transgenes or endogenous genes in ventral midbrain sequencing dataset. Fourth column indicates relative expression of transgenes and common marker genes. To control for differences in abundance of different cell types, values denote the mean percentage of reads +/-standard deviation corresponding to a given marker or transgene in cells positive for that marker or transgene.

| Transgenes | Number of expressing cells | Number of reads | Abundance in expressing cells |
| --- | --- | --- | --- |
| *tdTomato* | 1,927 | 7,354 | 0.15 +/- 0.31 |
| *FLPo* | 575 | 1,276 | 0.10 +/- 0.12 |
| *mCherry-IRES-Cre* | 299 | 639 | 0.11 +/- 0.13 |
| **Endogenous Genes** |  |  |  |
| *Snap25* | 8,053 | 19,143 | 0.08 +/- 0.05 |
| *Rbfox3* | 5,362 | 8,717 | 0.07 +/- 0.05 |
| *Slc17a6* | 1,648 | 2,249 | 0.05 +/- 0.04 |
| *Camk2a* | 3,137 | 4,521 | 0.06 +/- 0.05 |
| *Gad1* | 1,431 | 1,899 | 0.05 +/- 0.04 |
| *Gad2* | 2,581 | 4,288 | 0.07 +/- 0.05 |
| *Mog* | 236 | 325 | 0.13 +/- 0.08 |
| *Flt1* | 15 | 31 | 0.24 +/- 0.3 |

Once again, we clustered and annotated all cell types and assign them to broad categories: neurons, astrocytes, oligodendrocytes, microglia, and endothelia. These analyses revealed that 12,314 of 13,412 (91.81%) of the FACS-sorted and profiled nuclei were neuronal (Fig. 5B, C). We then separately subclustered excitatory and inhibitory neurons. This analysis yielded 7,019 excitatory (*Slc17a6*^+^) and 5,295 inhibitory (*Gad1^+^/Gad2^+^*) neurons (Fig. 5C). We focused subsequent analyses on inhibitory populations, overlaying viral transgene expression on the inhibitory clusters (Fig. 5D, E).

The *mCherry-IRES-Cre* and *FLPo* were present in several populations (Fig. 5E). The *mCherry-IRES-Cre* was notably abundant in a population that expressed markers such as *Pax6, Cdh23,* and *Pde11a* (Fig. 5E). This was intriguing because *Pax6* and *Cdh23* have been reported to be expressed in ZI, particularly in the ventral subdivision in which expression of GABAergic markers is dense, and the projection from ZI to VM has been shown to be GABAergic (Barthó et al., 2002b; Watson et al., 2014). For this reason, we further pursued this population. We injected *AAVrg-Cre* into VM and used RNAscope to measure *Pax6* co-expression in *Cre*-expressing cells (Fig. 6). Of 173 *Cre*^+^ cells in ZI, 123 (71%, n = 3 animals) co-expressed *Pax6,* confirming that it is a marker for this projection population. Thus, VECTORseq identified a novel subcortical inhibitory projection population.

**Figure 6.**
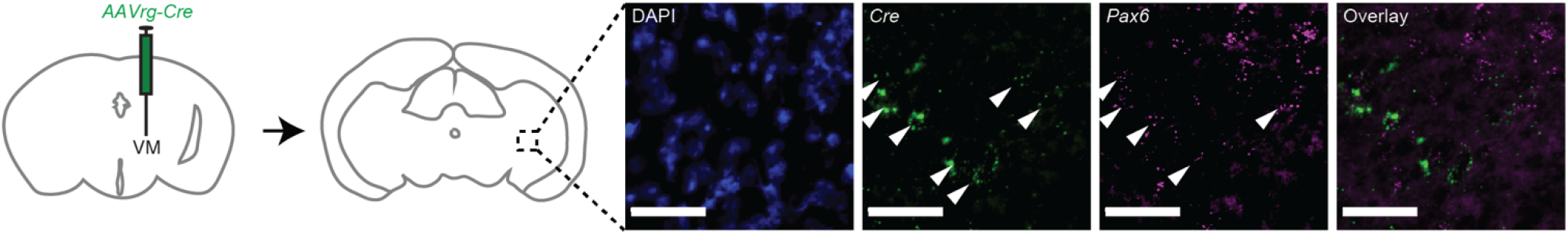
Analysis of candidate marker gene expression in ZI->VM population. **A.** Left, schematic of experiment. *AAVrg-Cre* was injected in right VM. 3 weeks later, mice were perfused and RNAscope was performed on ZI. Right, representative images. Middle left image shows expression of *Cre*. Arrowheads indicate *Cre^+^* cells. Middle right image shows *Pax6* expression. Arrowheads indicate *Cre^+^* cells that are also *Pax6*^+^. Scale bars, 50 μm.

To determine whether viral infection perturbed endogenous gene expression in these neurons, we then performed within-cluster comparisons of transgene^+^ and transgene^-^ cells in clusters. In two of the clusters analyzed, transgene^+^ cells were interspersed throughout, suggesting that the viruses do not perturb endogenous gene expression, as we had observed in SC (Supplementary Fig. 5A-D). Interestingly, in the smallest population analyzed, the *Pax6*-expressing cluster 10, transgene^+^ cells were interspersed with transgene^-^ cells but found on one side of the cluster (Supplementary Fig. 5E). After subclustering, nearly all transgene^+^ cells fell into one of the subclusters, where they were interspersed with many transgene^-^ cells (Supplementary Fig. 5F). This suggested that cluster 10 may correspond to two closely related *Pax6*^+^ populations that were not distinguished during our initial clustering, perhaps due to the small number of cells or because they differed relatively subtly in gene expression (Supplementary Fig. 5E, F). Further analyses revealed several endogenous neuronal genes whose expression differed between these two populations, including *Gfra1*, *Unc13c*, and *Ephb1*, suggesting that the projection from ZI to VM corresponds to a subtype of *Pax6*^+^ cells (Supplementary Fig. 5G-J). Taken together, these analyses suggest both that the retrograde viral transgenes did not perturb endogenous gene expression and that their distribution within clusters may be useful to guide subclustering of closely related subtypes.

## Discussion

The brain contains myriad projection neurons whose molecular and functional properties are unknown. Existing methods for transcriptionally profiling single projection neurons have been fruitful but are slow, laborious, costly to implement, and involved, requiring specialized equipment and separate processing and sequencing for each population. In theory, different populations could be pooled for a single sequencing run to reduce costs, but attempting to pool samples with methods such as cell hashing, which are not widely used, would entail additional steps and costs, and can reduce both yield and data quality (Gaublomme et al., 2019; Stoeckius et al., 2018). Another recently described method, BARseq, requires costly specialized equipment to perform *in situ* sequencing, precise calibration of viral expression to avoid toxicity and perturbation of gene expression, and prior knowledge of markers for cell types in the tissue of interest which, in the only published application, were obtained through conventional single-cell sequencing. In contrast, we developed and validated a new method, VECTORseq, that enables a theoretically limitless number of projection populations to be barcoded simultaneously and identified without the need to optimize additional steps or specialized equipment such as an intracellular recording system or flow cytometer (although the latter is compatible with VECTORseq, if desired). The isolated cells and nuclei can be sequenced in one run, rather than separate sequencing reactions for each projection target, reducing costs and increasing scalability. In comparison with existing approaches, VECTORseq is straightforward to implement and greatly reduces the number of animals sacrificed, sequencing costs, time, and steps (and potential failure points) required to characterize projection populations.

Using VECTORseq, we were able to detect a variety of functionally different transgenes delivered by commonly used retrograde viruses such as AAVrg and HSV under the control of several promoters, including *Synapsin*, *CAG*, and *EF1*α; surprisingly, we were also able to robustly detect viruses delivered retrogradely by AAV1, which is known to infect retrogradely but much less efficiently than AAVrg or HSV (Tervo et al., 2016). Thus, VECTORseq is a highly sensitive method that we expect will be compatible with any viruses (such as Cav-2, lentivirus, and others) that are used to target projection types (Wickersham and Feinberg, 2012).

Importantly, comparison of virally infected and uninfected cells within clusters showed no differences in the expression of endogenous genes, and the markers for virally labeled clusters did not appear enriched for inflammatory or antiviral genes, suggesting that both AAV and HSV did not interfere with the “ground state” gene expression of these cell types. We combined as many as 6 different viruses and targeted up to 3 structures in individual proof-of-principle experiments, but future experiments could label a vast array of projection targets due to the diversity of available transgenes (such as recombinases, fluorescent proteins, optogenetic and chemogenetic tools) and ability to distinguish closely related sequences such as different *Cre* variants. In order to study structures harboring a large number of projection populations, such as SNr, which was recently shown to innervate 39 different targets (McElvain et al., 2021), it would be possible to further increase labeling diversity through generation of custom AAV containing noncoding barcode sequences.

We performed 5’ sequencing in this study because many viruses share a 3’ UTR element (WPRE) that boosts transgene expression (Wang et al., 2016). Although most neuronal studies use 3’ sequencing, the core facilities we contacted are able to perform 3’ or 5’ sequencing using commercial kits at identical costs and without additional steps for the end user. It may be possible to perform 3’ sequencing with VECTORseq by injecting viruses that lack the WPRE; however, one of our viruses*, hSyn-Dre*, lacked a WPRE, and only one read was detected for this virus, suggesting that the WPRE may increase RNA expression or stability and thus detectability in sequencing (Wang et al., 2016). One alternative approach could be to perform 3’ sequencing and include a primer targeted to the WPRE in order to amplify the unique viral transgene sequences upstream of the WPRE. Another alternative to 5’ sequencing would be to use single-cell platforms that sequence through gene bodies; for example, in preliminary studies we were able to detect viral transgenes using Smart-Seq2 (data not shown) (Picelli et al., 2014).

We first validated VECTORseq by labeling visual cortical projection populations, finding that viral barcodes were in excitatory neurons, as expected. We then combined validation and discovery by investigating structures with a mixture of known and unknown projections. First, VECTORseq identified a known projection population in superior colliculus (SC) that innervates contralateral PPRF and expresses *Pitx2,* uncovering an additional marker for this population, *Pmfbp1* (Fig. 7A) (Masullo et al., 2019; Xie et al., 2021). Second, we examined populations that project to the thalamic nucleus LP, the rodent homolog of pulvinar. It is well known that wide-field cells in superficial SC project to the lateral portion of LP, and a fortuitously identified Cre transgenic line (that does not recapitulate endogenous gene expression) has been widely adopted in the last few years for functional studies of WF cells and LP physiology (Gale and Murphy, 2014, 2016; Hoy et al., 2019; Reinhard et al., 2019; Sans-Dublanc et al., 2021).

**Figure 7.**
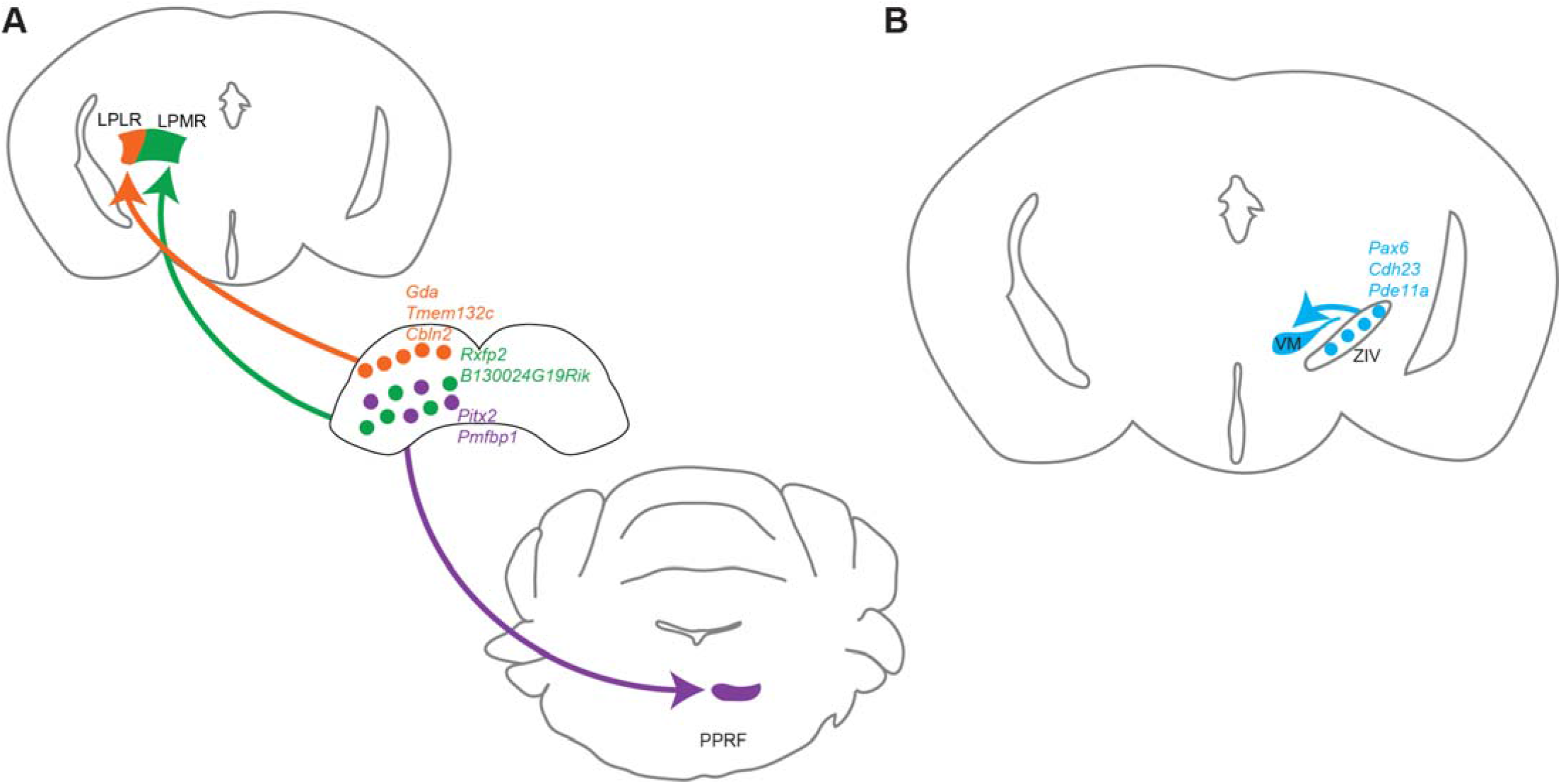
Summary of cell types and markers identified using VECTORseq in this study. **A.** VECTORseq identified new markers for SC WF cells (including *Gda, Tmem132c*, and *Cbln2*) that project to LPLR and for *Pitx2*^+^ cells that project to PPRF (*Pmfbp1*). In addition. VECTORseq identified a hitherto elusive population of deep SC neurons that project to LPMR labeled by *Rxfp2* and *B130024G19Rik*. **B**. VECTORseq identified ZI *Pax6*^+^ cells as GABAergic neurons that project to thalamic nucleus VM. These cells also express *Cdh23* and *Pde11a*.

However, the molecular identity of WF cells was previously unknown. Using VECTORseq, we were able to define WF cells using SC transcriptional profiling (Fig. 7A). Interestingly, a recent study used Patch-seq to transcriptionally profile this population, also finding that *Cbln2* and *Gda* are expressed in LP-projecting superficial SC neurons that appear to correspond to WF cells (Xie et al., 2021). Thus, our data match closely with those obtained using much more labor-intensive and less scalable established methods, providing further validation of VECTORseq. In addition to these known populations, we identified a long-elusive population: For more than forty-five years it has been known that the medial portion of pulvinar, the homolog of LPMR, receives input from the deep, oculomotor layers of SC, which may explain why LP neurons are modulated by movements, yet the properties of this SC projection population have remained mysterious (Benevento and Fallon, 1975). Anterograde tracing from mouse SC has revealed similar projections to LPMR but the functional and molecular properties of this population have never been defined in any species (to our knowledge) (Gharaei et al., 2020). Using VECTORseq, we were able to identify this deep SC population that projects to LPMR and specific markers for it, including *Rxfp2* and the highly expressed uncharacterized gene *B130024G19Rik*. Just as the discovery of a Cre transgenic line that labels WF cells has been transformative, the identification of these markers could enable functional studies of this elusive LP-projecting SC population and its role in sensory processing, sensorimotor integration, and behavior (Fig. 7A). In this way, VECTORseq both enabled molecular characterization of defined populations and discovery of previously undefined populations.

Finally, to determine its applicability to inhibitory projection neurons, we performed VECTORseq on ventral midbrain neurons that project to several motor structures. This analysis defined a GABAergic population in zona incerta that expresses the marker *Pax6* and innervates VM (Fig. 7B). Interestingly, one previous study found that GABAergic neurons in ZI project to VM, while a separate study analyzing gene expression suggested that *Pax6* was found in the ventral portion of ZI where GABAergic cells are enriched (Barthó et al., 2002a; Watson et al., 2014). Here, we used an unbiased approach to link these disparate observations, illustrating the power of VECTORseq to solve correspondence problems and relate gene expression to connectivity and functional properties.

### Limitations

Cav-2 was not detected in our ventral midbrain sequencing dataset, but this was because the injection failed, illustrating that, as for any retrograde approach, success hinges on infection efficiency. It may be possible to increase the efficiency of Cav-2 labeling through heterologous expression of its receptor, *hCar*, in the source structure of interest (Li et al., 2018). Conversely, a second caveat is that our attempts to sequence projections to a structure very close to the source structure (SC to CnF) appear to have been confounded by ambient RNA from the injection site. Therefore, when examining projection targets near the source structure, it may be preferable to ensure that dissections exclude the injection site or to use a retrograde virus that does not yield strong expression at the injection site, such as HSV, which has been reported to express long-term in retrogradely labeled neurons but only transiently at the injection site (Fenno et al., 2014).

### Concluding remarks

In conclusion, we devised a new technology, VECTORseq, to address a major technical challenge in the field: relating the molecular identities of intermingled neuronal populations with their connectivity, functional properties, and behavioral functions. In contrast to existing methods, VECTORseq is straightforward to implement, requires no specialized equipment, uses commercially available components, and enables multiplexing to reduce the number of animals sacrificed and sequencing reactions performed. We rigorously demonstrated that transgenes delivered by retrogradely infecting viruses are detected and discriminated in single-cell sequencing data. We then validated VECTORseq in several brain areas and myriad projection populations, identifying molecular markers that could be used for targeted monitoring and manipulation of cell types and to understand their physiology. In the future, VECTORseq could be adapted to incorporate other viral tracers, including those that traffic anterogradely. Our study thus establishes a simple yet powerful approach to link the transcriptional identities of neuronal cell types to their connectivity and delineates new subcortical cell types involved in sensorimotor integration.

## Acknowledgments

We thank S. Darmanis, Z. Knight, E. Macosko, B. Wu, and members of the Feinberg laboratory for helpful discussions and comments on the manuscript. We thank C. Cheung for help with software development and code refactorization. We thank J. McGuire and M. Bernardi at the Gladstone genomics core for 10X library preparation and sequencing and staff at the UCSF Center for Advanced Technology (CAT) core for advice and support with RNA sequencing. Flow cytometry was performed by the Gladstone flow cytometry core, which is supported by NIH S10 RR028962, James B. Pendleton Charitable Trust, and NIH P30 AI027763. This work was supported by departmental funds and grants from the E. M. Ziegler Foundation for the Blind, Sandler Foundation, Klingenstein-Simons Fellowship Award in Neuroscience, Brain and Behavior Research Foundation (NARSAD Young Investigator Awards 25337 and 27320), Whitehall Foundation, Simons Foundation (SFARI 574347), and US National Institutes of Health (DP2 MH119426 and R01 NS109060) to E.H.F.

## Author contributions

Victoria Cheung: Methodology, Software, Validation, Formal Analysis, Investigation, Resources, Data Curation, Writing—original draft, Writing—Review & Editing, Visualization, Project Administration; Philip Chung: Software, Validation, Formal analysis, Writing—Review & Editing, Visualization; Yesenia Colleen Lopez: Investigation (pilot experiments), Writing—Review & Editing; Max Bjorni: Investigation, Writing—Review & Editing; Varvara Alexeevna Shvareva: Investigation, Writing—Review & Editing; Evan H. Feinberg: Conceptualization, Methodology, Resources, Writing—original draft, Writing—Review & Editing, Visualization, Project Administration, Funding Acquisition, Supervision

---

**Supplementary Figure 1:**
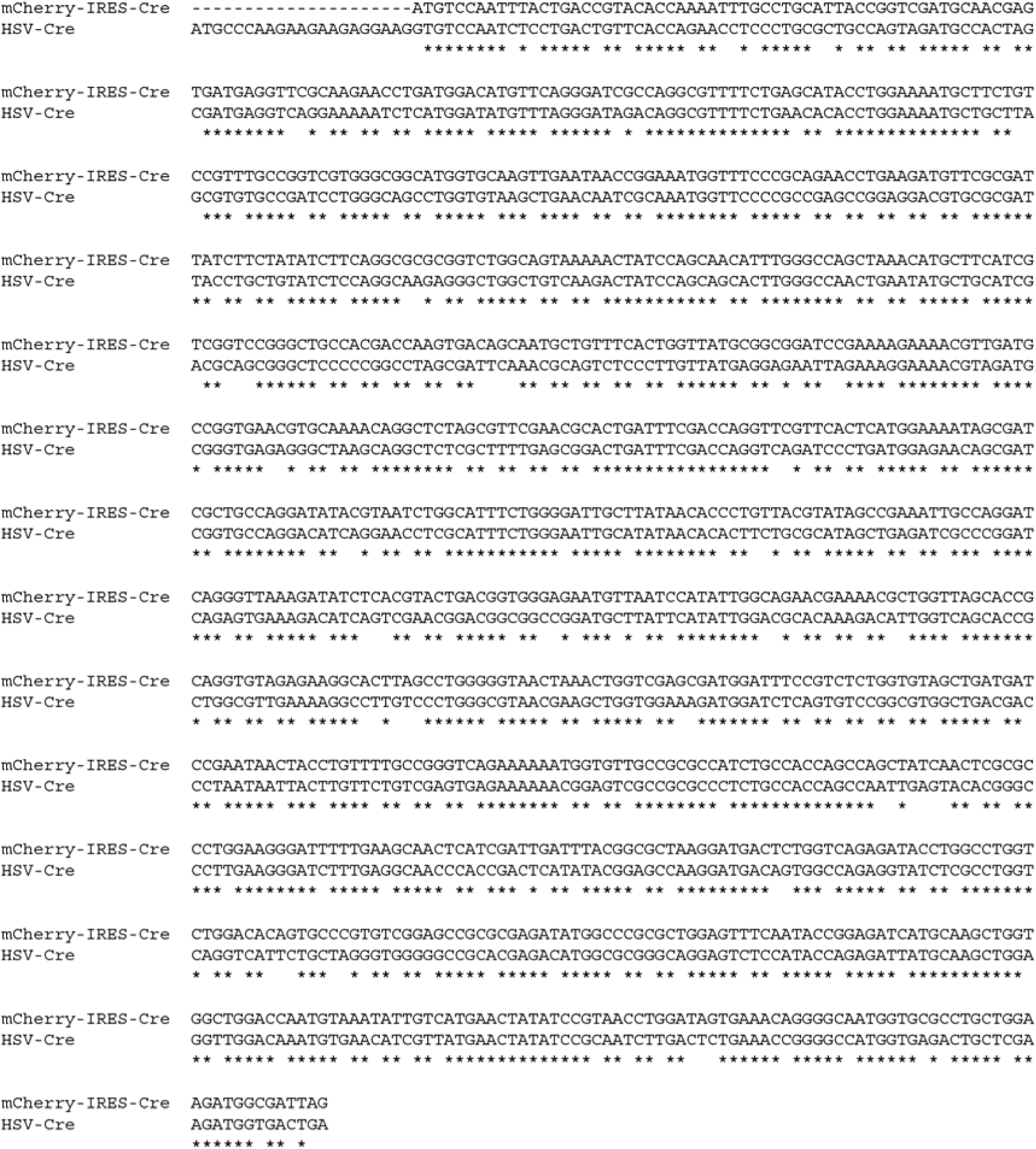
Alignment of *Cre* coding sequences used in the study. Alignments were performed using ClustalW. Asterisks indicate identities and gaps indicate mismatches. The *Cre* sequence in *mCherry-IRES-Cre* contains a 5’ SV40 nuclear localization signal.

**Supplementary Figure 2.**
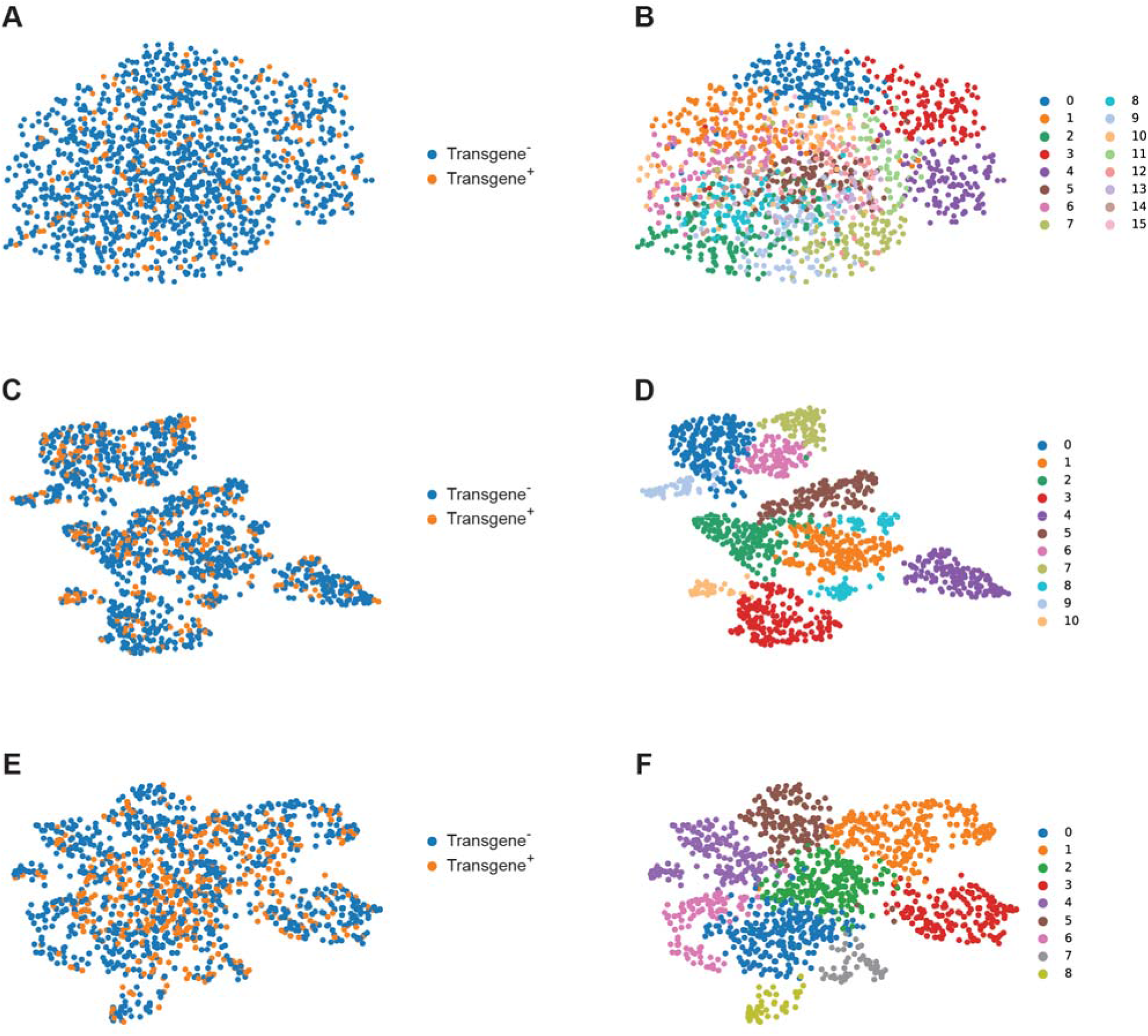
Analysis of the effects of viral transgene expression on endogenous gene expression within clusters in SC. Transgene^+^ and transgene^-^ cells are interspersed in clusters 7 (**A**), 10 (**C**), and 11 (**E**) from Fig. 3. Transgene^+^ and transgene^-^ cells remain interspersed after further subclustering these clusters (**B, D, F**), indicating that AAV and HSV do not perturb endogenous gene expression in these cells.

**Supplementary Figure 3.**
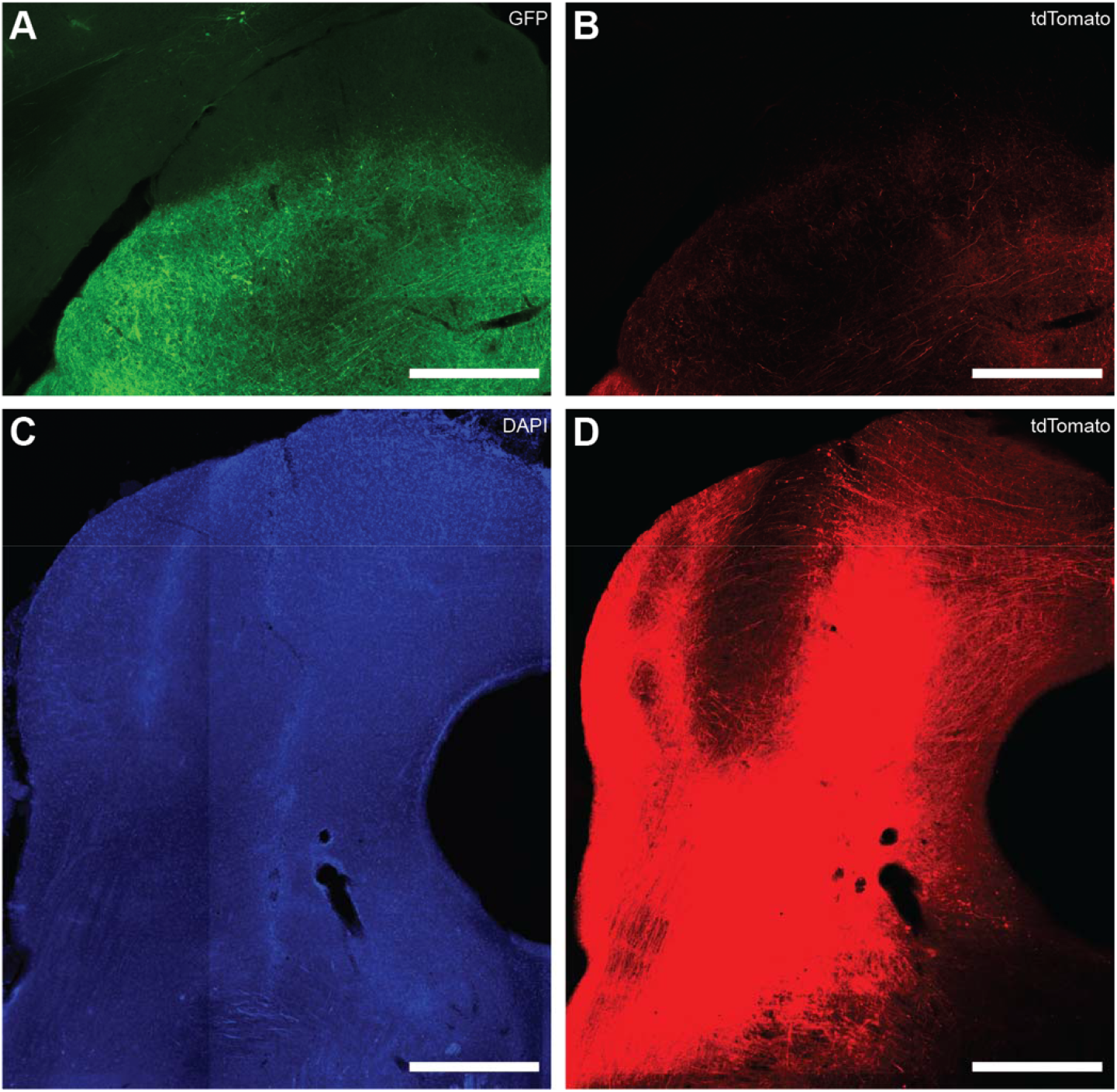
Absence of retrograde labeling from CnF in SC. **(A)** Retrogradely infected SC neurons from *AAVrg-CAG-GFP* injection into contralateral PPRF. **(B)** Only sparse tdTomato^+^ fibers and no tdTomato^+^ cell bodies are seen in SC from injection of *AAVrg-CAG-tdTomato* in left CnF. **(C, D)** Area around injection site in CnF shows intense tdTomato labeling due to local infection. 50 μm tissue section in A and B taken from roughly 4.0 mm posterior to bregma. 50 μm tissue section in C and D taken from roughly 4.95 mm posterior to bregma. SC extends caudally to roughly 4.85 mm posterior to bregma. Consequently, 300 μm sections collected for SC tissue microdissection included portions of injection site. Images in A, B, and D were collected with the same confocal settings and processed identically to allow comparisons of brightness.

**Supplementary Figure 4.**
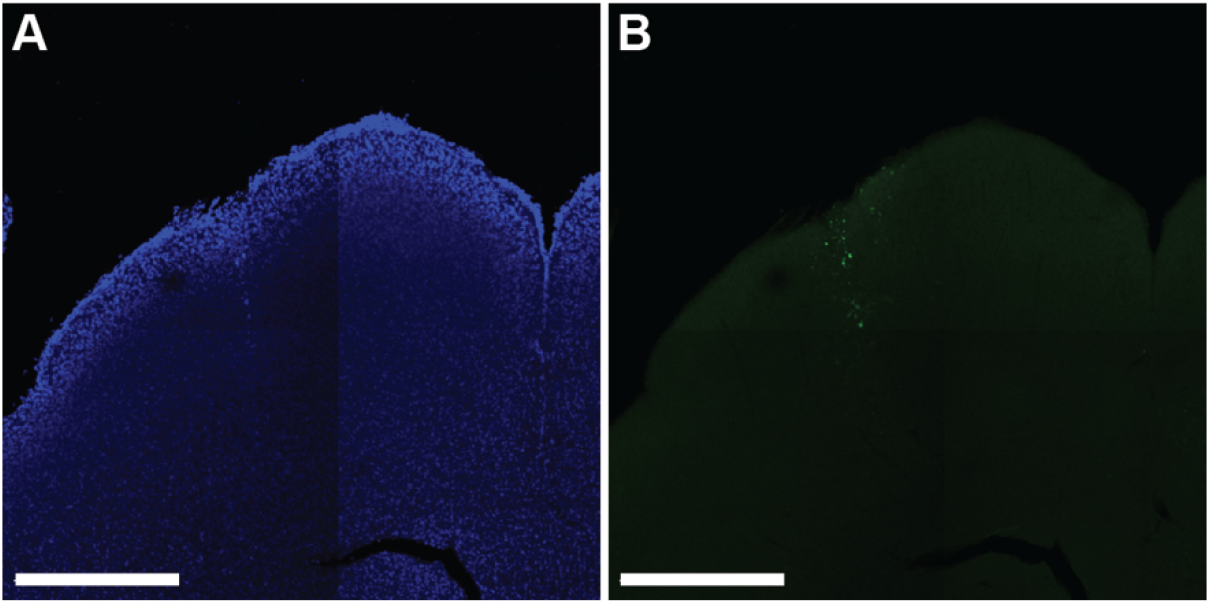
*Cav2-GFP* injection site in SC. **(A)** DAPI labeling in SC. Injection track is faintly visible as nearly vertical line of bright labeling. **(B)** GFP labeling in SC. Only a few cells are labeled and they localize to the injection track.

**Supplementary Figure 5.**
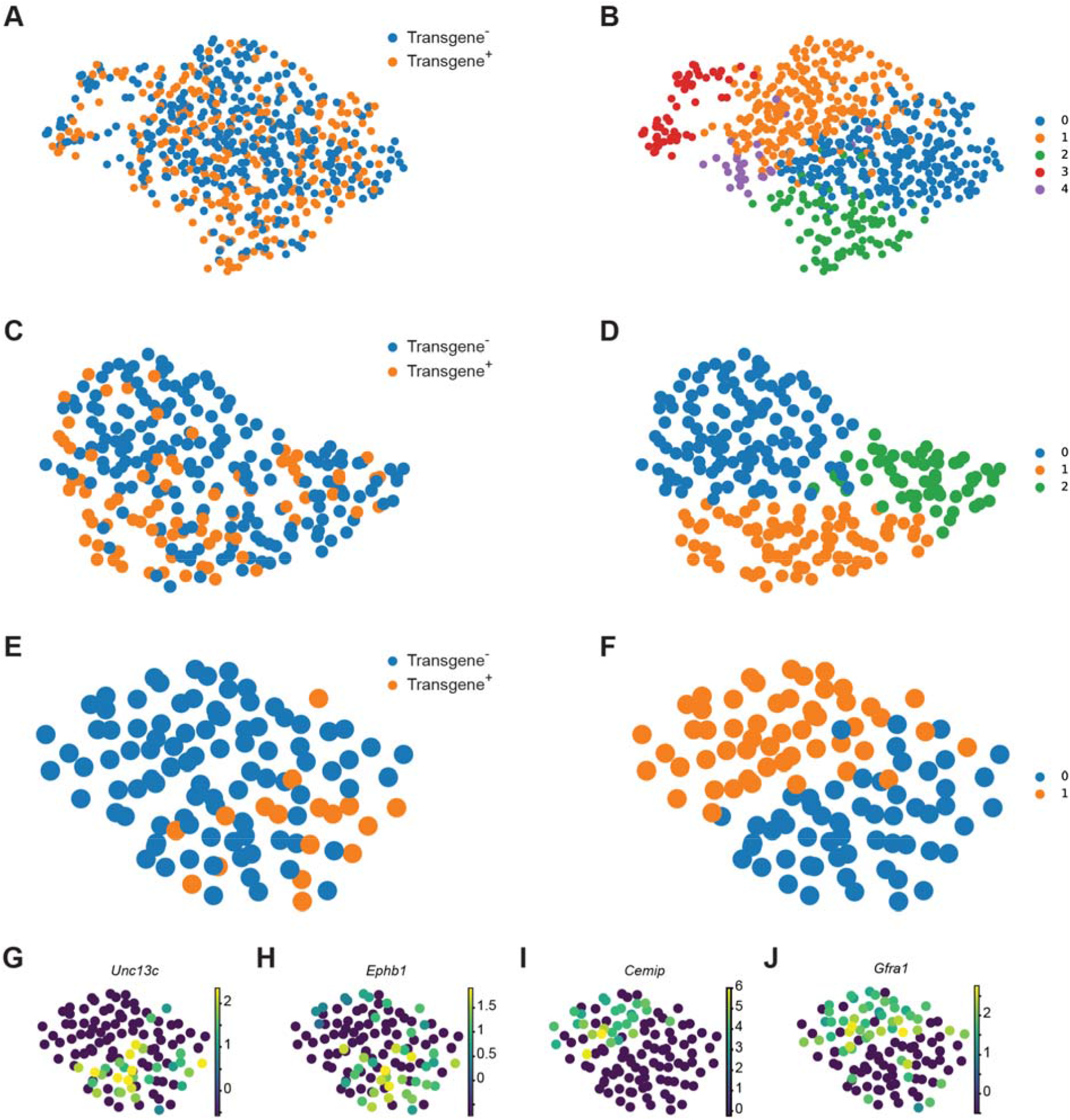
Analysis of the effects of viral transgene expression on endogenous gene expression within clusters in ventral midbrain. Transgene^+^ and transgene^-^ cells are interspersed in clusters 2 (**A**), 5 (**C**), and 10 (**E**) from Fig. 5. Transgene^+^ and transgene^-^ cells remain interspersed in subclusters derived from clusters 2 and 5 clusters (**B, D**), indicating that AAV does not perturb endogenous gene expression in these cells. Transgene^+^ cells are interspersed with transgene^-^ cells in cluster 10, indicating that the transgenes did not disrupt viral transgene expression, but appear enriched in one of two subclusters (**F**), suggesting that this cluster may correspond to two closely related subtypes. (**G-L**) Genes differentially expressed in the two subclusters of cluster 10. Scale bars indicate relative expression.

## Materials and Methods

### Stereotaxic surgeries

All experiments were performed according to Institutional Animal Care and Use Committee standard procedures. All surgeries were performed on adult (8-12 weeks) male C57BL/6J mice from the Jackson Laboratory (JAX, stock 000664). Each set of histology surgeries used between 4-5 mice. For each set of sequencing experiments, between 4-6 mice were used. Mice were administered buprenorphine 30 minutes prior to anesthesia. 30 minutes later, mice were anesthetized with isoflurane and given meloxicam for analgesia. All coordinates are in mm. Angled injections were always done such that the tip of the syringe pointed towards the midline and the plunger tilted away from the midline. Injection coordinates were determined using an adult mouse atlas.

#### Viruses used

For sequencing:

*AAVrg-CAG-GFP*, Addgene
*AAVrg-CAG-tdTomato*, Addgene
*AAVrg-Ef1α-mCherry-IRES-Cre*, Addgene
*AAVrg-Ef1α-FLPo*, Addgene
*AAVrg-hSyn-Dre*, Addgene
*AAVrg-hSyn-TurboRFP*, Addgene
*HSV-hEF1α-Cre (RN425)*, MGH Gene Delivery Technology Core
*Cav2-GFP*, IGMM

For *in situ* hybridization experiments using RNAscope:

*AAVrg-hSyn-Cre*, Addgene
*AAV1-CAG-FLEX-tdTomato*, Addgene

#### Injection coordinates and volumes (all measurements in mm)

**V1**, 35 nl/depth at a rate of 10 nl/minute at the coordinates:

1.) AP: 2.69 posterior to bregma, ML: 2.50, DV: 0.60, 0.40 below pia
2.) AP: 2.91 posterior to bregma, ML: 2.50, DV: 0.60, 0.40 below pia
3.) AP: 3.15 posterior to bregma, ML: 2.50, DV: 0.75, 0.50 below pia
4.) AP: 3.30 posterior to bregma, ML: 2.50, DV: 0.50, 0.25 below pia
5.) AP: 3.51 posterior to bregma, ML: 2.50, DV: 0.50, 0.25 below pia
6.) AP: 3.79 posterior to bregma, ML: 2.50, DV: 0.50, 0.25 below pia
7.) AP: 4.00 posterior to bregma, ML: 2.50, DV: 0.50, 0.25 below pia
8.) AP: 4.25 posterior to bregma, ML: 2.50, DV: 0.55, 0.40 below pia

**Striatum**, injection rate of 30nl/minute at the coordinates:

1.) AP: 0.90 anterior to bregma, ML: 1.50, DV: 2.00 below pia, 150 nl at single depth
2.) AP: 0.45 anterior to bregma, ML: 2.00 ML, DV: 2.00 below pia, 150 nl at single depth
3.) AP: 0.00 at bregma, ML: 2.25, DV: 2.50 below pia, 150 nl at single depth
4.) AP: 0.34 posterior to bregma, ML: 2.50, DV: 2.50 below pia, 150 nl at single depth

**SC**, 50 nl/depth at a rate of 30 nl/minute at the coordinates:

1.) AP: 0.25 anterior to lambda, ML: 1.00, DV: 2.00, 1.75, 1.50, 1.25, 1.00 below skull surface
2.) AP: 0.50 anterior to lambda, ML: 1.00, DV: 2.00, 1.75, 1.50, 1.25, 1.00 below skull surface

**Ventral midbrain**, 100 nl/depth at a rate of 30 nl/depth, 10° angle at the coordinates:

1.) AP: 1.25 anterior to lambda, ML: 2.28, DV: 4.60, 4.40, 4.20 below skull surface

**VM,** injection rate of 30 nl/minute at the coordinates:

1.) AP: 1.23 posterior to bregma, ML: 0.75, DV: 4.15 below skull surface, 150 nl at single depth
2.) AP: 1.43 posterior to bregma, ML: 1.00, DV: 4.25 below skull surface, 50 nl at single depth
3.) AP: 1.67 posterior to bregma, ML: 0.75, DV: 4.25 below skull surface, 50 nl at single depth

**MLR**, injection rate of 30 nl/minute at the coordinates:

1.) AP: 4.23 posterior to bregma, ML: 1.30, DV: 3.83 below skull surface, 70 nl at single depth
2.) AP: 4.43 posterior to bregma, ML: 1.25, DV: 3.63 below skull surface, 70 nl at single depth
3.) AP: 4.63 posterior to bregma, ML: 1.20, DV: 3.80, 3.40 below skull surface, 30 nl/depth
4.) AP: 4.83 posterior to bregma, ML: 1.50, DV: 3.50 below skull surface, 50 nl at single depth
5.) AP: 4.89 posterior to bregma, ML: 1.00, DV: 3.00 below skull surface, 100 nl at single depth

**PPRF**, injection rate of 30 nl/minute at the coordinates:

1.) AP: 4.95 posterior to bregma, ML: 0.63, DV: 5.13, 4.88, 4.63, and 4.38 below skull surface, 50 nl per depth
2.) AP: 5.07 posterior to bregma, ML: 0.50, DV: 5.13, 4.88, 4.63, and 4.38 below skull surface, 50 nl per depth
3.) AP: 5.19 posterior to bregma, ML: 0.50, DV: 5.13, 4.88, 4.63, and 4.50 below skull surface, 50 nl per depth
4.) AP: 5.33 posterior to bregma, ML: 0.50, DV: 5.33, 5.25, 5.00, and 4.75 below skull surface, 50 nl per depth

**CnF**, 50 nl/depth at an injection rate of 30 nl/minute at the coordinates:

1.) AP: 4.83 posterior to bregma, ML: 1.13, DV: 2.85 below skull surface
2.) AP: 4.95 posterior to bregma, ML: 1.13, DV: 2.85 below skull surface
3.) AP: 5.07 posterior to bregma, ML: 1.37, DV: 3.13, 2.85 below skull surface
4.) AP: 5.19 posterior to bregma, ML: 1.25, DV: 2.85 below skull surface

**LP**, 70 nl/depth at an injection rate of 30 nl/minute at the coordinates:

1.) AP: 1.55 posterior to bregma, ML: 1.00, DV: 2.63 below skull surface
2.) AP: 1.67 posterior to bregma, ML: 1.00, DV: 2.63 below skull surface
3.) AP: 1.79 posterior to bregma, ML: 1.50, DV: 2.63 below skull surface
4.) AP: 1.91 posterior to bregma, ML: 1.37, DV: 2.63 below skull surface
5.) AP: 2.03 posterior to bregma, ML: 1.30, DV: 2.50 below skull surface
6.) AP: 2.15 posterior to bregma, ML: 1.25, DV: 2.60 below skull surface
7.) AP: 2.27 posterior to bregma, ML: 1.37, DV: 2.60 below skull surface

### Single Cell Isolation

#### Tissue preparation

Mice were anesthetized and trans-cardially perfused with 4 °C aCSF. Brains were quickly dissected out and placed in chilled slurry of N-methyl-d-glucamine Buffer (NMDG Buffer). Brains were then glued in the coronal orientation onto a vibratome platform. The vibratome was filled with the cold NMDG buffer slurry. 300 micrometer sections were sliced at 0.06 mm/second and sections with the region of interest (ROI) were isolated. The slices were further micro-dissected to isolate the ROI. The ROI were recovered in a 37°C NMDG bath for 20 minutes before being placed in room temperature aCSF for 20 minutes.

#### Single cell dissociation

Tissue was processed using the Papain Dissociation System Protocol (Worthington Biochemical Corporation, LK003150). In summary, this protocol involved gentle trituration using a transfer pipette (Falcon, 357524) every 20 minutes for 1-1.5 hours using the provided dissociation buffers in a 37°C rocker. After dissociation, the suspension was centrifuged in a low-bind microcentrifuge tube (Eppendorf) at 300 *g* for 10 minutes. After centrifugation and removal of supernatant, the pellet was resuspended with provided albumin-ovomucoid inhibitor. Cell debris was removed using the provided density gradient solutions, spinning in a centrifuge at 100 *g* for 7 minutes. Supernatant was discarded; cell pellet was reconstituted in 1mL of aCSF. Cells were counted using a hemocytometer and diluted or concentrated to roughly 450 cells/μL. 10x Genomics 5’ v1.1 library prep and NextSeq sequencing were performed by the Gladstone Institute Genomics core.

### Single Nuclei Isolation

#### Tissue preparation

Mice were anesthetized and trans-cardially perfused with 4 °C aCSF. Brains were quickly dissected out and placed in clean 4 °C aCSF. Brains were then glued in the coronal orientation onto a vibratome platform. The vibratome was filled with 4 °C aCSF. 300 micrometer sections were cut at 0.06 mm/second and sections containing the ROI were isolated. The ROI was micro-dissected out of the brain slices, diced into rice-sized pieces, and then placed in a low-bind microcentrifuge tube (Eppendorf). The tissue was then flash-frozen using liquid nitrogen.

#### Single Nuclei dissociation

We followed a published protocol (Martin et al., 2020). Briefly, all steps were done either in 4° C or on ice. All reagents and items used were pre-chilled overnight at 4° C. Flash frozen tissue was gently triturated with detergent-based extraction buffer until tissue was visibly broken up, careful not to generate any bubbles. The entire volume was then passed through a 26G needle twice before transfer into a pre-chilled 50mL Falcon tube. 30 mL of wash buffer (HEPES-based buffer with 10% BSA) was added. This volume was then split into 2 15 mL centrifuge tubes for centrifugation at 600g for 10 minutes at 4° C. Supernatant was removed until roughly 500 μL remained in each tube. Samples were then pooled together (total volume = 1 mL). The suspension was then passed through a pre-chilled 40 micrometer cell strainer and filtered using only gravity. Nuclei were counted using a hemocytometer and diluted or concentrated to roughly 8-10 million nuclei/mL. 200 μL of nuclei were reserved as a negative control. The remaining volume of nuclei was stained with rabbit anti-NeuN antibody conjugated to AlexaFluor488 (Abcam, ab190195) at a concentration of 0.1-10 micrograms/mL for 30 minutes in the dark on a gentle rocker. Stained nuclei were then washed with FACS buffer, centrifuged at 200 *g* for 1 minute, and supernatant was aspirated. The pellet was then resuspended in 1 mL of FACS buffer. DAPI was then added at 1 μg/μL. FACS was performed at the Gladstone Institute Flow Core on an Aria II. 10x Genomics 5’ v1.1 and v2 library prep was performed by the Gladstone Institute Genomics core. Sequencing of the library was done with UCSF’s CAT core.

### Sequencing Analysis

Analyses were performed in Python. 10x Genomics’ Cellranger cli was used to add viral transgenes to their mouse (mm10) reference genome (Zheng et al., 2017). Fastq outputs were aligned to this customized mm10 reference genome; introns were included in the alignment. Outputs of the alignments include a count matrix. Using Scanpy, cells with more than 5% mitochondrial gene expression were filtered out and counts in each cell were normalized to 10^4^ and then log-normalized (Wolf et al., 2018). We implemented a version of term frequency-inverse document frequency (TF-IDF) normalization using the formula:

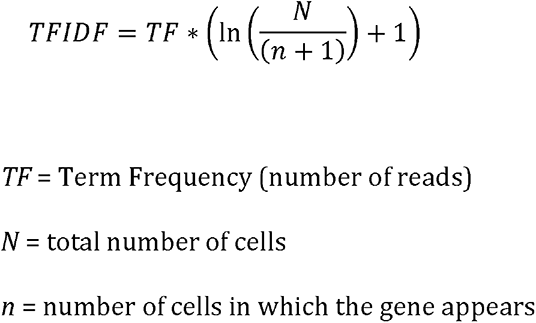

TF-IDF is useful for weighting genes according to their variance across the population rather than their absolute expression (Moussa and Măndoiu, 2018). After normalization steps, the top 2000 highly variable genes were selected. Data were subset using these highly variable genes. The subset data were then scaled to unit variance and zero-centered. Data preprocessing in Scanpy involved dimensionality reduction using principal component analysis (PCA) with 50 principle components and *svd_solver* set to “*arpack*”, followed by constructing a neighborhood graph using 15 nearest neighbors. Only endogenous genes were used to create the neighborhood graph to avoid any potential influence of viral transgenes on clustering. The Leiden algorithm was applied to the neighborhood graph with resolution 0.6 to generate clusters. Uniform manifold approximation projection (UMAP) was applied to the neighborhood graph to visualize the resultant clusters in 2 dimensions. Dendrogram plots were generated using complete-linkage hierarchical clustering using Pearson’s correlation coefficient and top 50 principal components.

### Marker Gene Selection

Top 50 genes from each cluster were selected using the Mann-Whitney *U* test with Benjamini-Hochberg procedure to control for false discovery rate. Gene expression heatmaps were generated of the top 50 differentially expressed genes from each cluster. Clusters were merged based on visual analysis of heatmaps, dendrogram plots, and applying an 80% cut-off to mutual presence of the top 50 unique genes across cluster pairs using the Jaccard similarity score.

After cluster merging, genes that are unique to a specific cluster, highly expressed, and expressed in the majority of that specific cluster were selected as marker genes. Subsets of these unique genes were selected as biomarkers of interest for *in situ* hybridization using RNAscope.

### Histology

#### Tissue preparation for native fluorescence

Mice were anesthetized with 100% isoflurane and transcardially perfused first with dPBS and subsequently with 10% formalin solution for fixation. Brains were then harvested and post-fixed in 10% formalin for 4-12 hours at 4°C. After post-fixation, brains were transferred to a 20% sucrose solution and kept at 4 °C until the tissue was saturated with sucrose and no longer floating in solution. Brains were then frozen in OCT (Sakura) and sectioned coronally via cryostat at a thickness of 50 micrometers.

#### Tissue preparation for *in situ* hybridization

Mice were rapidly anesthetized with 100% isoflurane and decapitated. Brains were quickly dissected and placed in OCT-filled cryo-sectioning cubes and immediately transferred into a slurry bath of 100% ethanol and dry ice for flash freezing. Frozen brains were stored at -80 °C until ready for use. Brains were cryo-sectioned at -16 °C. Each section was 15 μm thick. Each section was directly mounted onto *Superfrost Plus* slides (Fisher) and dried for at least 30 minutes inside the cryostat chamber before storage at -80 °C.

#### RNAscope

All experiments were performed according to the Advanced Cell Diagnostics (ACD) RNAscope protocol. Each Target Probe contains a mixture designed to bind to a specific target RNA. Each of these probes were detectable in one of three color channels, C1, C2, and C3 as follows: C1, Alexa 488 nm; C2, Atto 550 nm; C3, Atto 647 nm. In each experiment, we probed for *Cre* in channel 1(C1), *tdTomato* in channel 2 (C2), and the gene of interest (*Gda, Pax6, Pitx2, Pmfbp1, Rxfp2*) in channel 3 (C3). Briefly, tissue was immediately fixed at 4°C in pre-chilled formalin for 15 minutes following removal from storage at -80°C. All dehydration and wash steps were performed at room temperature using 50% ethanol, 70% ethanol, and 100% ethanol. Protease IV pretreat (from the RNAscope Fluorescent Assay v1 kit) was used to permeabilize the tissue. Incubation occurred at room temperature for no more than 30 minutes to prevent over-digestion. After pretreatment incubation, slides were washed twice in PBS with gentle agitation for 30 seconds. The probe mix was applied to each slide and incubated for 2 hours at 40 °C. After probe incubation, slides were washed twice for 2 minutes each in RNAscope Buffer at room temperature. Probe hybridization signals were augmented using sequential hybridization of 4 amplifiers. Incubation times varied by amplification step but were all performed at 40 °C. Between each amplification incubation step, slides were washed twice for 2 minutes each in RNAscope buffer at room temperature. Amp4 Alt B-FL was used for the last amplification step. After the last wash, slides were coverslipped and imaged. RNAscope images in figures are pseudocolored for accessibility.

